# Notch effector Hes1 marks an early perichondrial population of skeletal progenitor cells at the onset of endochondral bone development

**DOI:** 10.1101/2020.03.13.990853

**Authors:** Yuki Matsushita, Mizuki Nagata, Joshua D. Welch, Sunny Y. Wong, Wanida Ono, Noriaki Ono

## Abstract

The perichondrium, a fibrous tissue surrounding the fetal cartilage, is an essential component of developing endochondral bones that provides a source of skeletal progenitor cells. However, perichondrial cells remain poorly characterized due to lack of knowledge on their cellular diversity and subset-specific mouse genetics tools. Single cell RNA-seq analyses reveal a contiguous nature of the fetal chondrocyte-perichondrial cell lineage that shares an overlapping set of marker genes. Subsequent cell-lineage analyses using multiple *creER* lines active in fetal perichondrial cells – *Hes1-creER*, *Dlx5-creER* – and chondrocytes – *Fgfr3-creER* – illustrate their distinctive contribution to endochondral bone development; postnatally, these cells contribute to the functionally distinct bone marrow stromal compartments. Particularly, Notch effector *Hes1-creER* marks an early skeletal progenitor cell population in the primordium that robustly populates multiple skeletal compartments. These findings support the concept that perichondrial cells participate in endochondral bone development through a distinct route, by providing a complementary source of skeletal progenitor cells.

## Introduction

The perichondrium is a poorly characterized fibrous tissue surrounding the fetal cartilage despite its essential roles in endochondral bone development – a highly organized process in which initial cartilage templates are replaced by bone and its marrow space. This process starts when chondrocytes and their surrounding perichondrial cells are defined from undifferentiated mesenchymal cells in condensations owing to actions of the transcription factor Sox9^1^. As chondrocytes organize the growth plate in their avascular environment, surrounding perichondrial cells develop as distinct cell types in their vascular-rich environment. The perichondrium provides two important functions; first, it provides cues for adjacent chondrocytes in the cartilage template^2^, and second, it provides a putative source of progenitor cells supporting skeletal development. Cellular heterogeneity and fates of perichondrial cells are not well defined.

Initially, the perichondrium is formed as elongated fibroblastic layers without differential morphological characteristics. When chondrocytes within the cartilage template undergo hypertrophy, the adjacent perichondrium becomes differentiated and forms the osteogenic perichondrium, partly due to actions of indian hedgehog (Ihh) released from pre-hypertrophic chondrocytes^3–5^. The osteogenic perichondrium is where the first osteoblast precursor cells expressing osterix (Osx) are formed; these cells can translocate into the marrow space and become osteoblasts^6^. However, these cells eventually disappear from the marrow space in the later stage of development^5,7^. The perichondrium persists well into the postnatal stage as long as the growth plate maintains its structure. Cathepsin K (Ctsk)-expressing cells in the perichondrial groove of Ranvier provide a source for other perichondrial and periosteal cells, which include a population of skeletal stem cells (SSCs) contributing to bone fracture healing^8–10^. In contrast, the cartilage template provides a principal cellular source of bone marrow stromal cells (BMSCs) including mesenchymal progenitor populations^11^; particularly, these precursor cells express Sox9^1,12^. However, how early perichondrial cells participate in endochondral bone development remains largely unclear.

Analyses of chondrocytes in early skeletal development have been facilitated by a well-defined set of tamoxifen-inducible mouse genetics tools, including *Col2a1-creER^13^*, *Sox9-creER^14^* and *Acan-creER^15^*; however, these lines can simultaneously mark perichondrial cells, at least to some extent^12^. Studies of perichondrial cells have been largely hampered due to lack of their subset-specific marker genes and mouse genetics tools; particularly, existing inducible genetic tools that are highly active in perichondrial cells, such as *Prrx1-creER^16^* and *Osx-creER^6^*, simultaneously mark chondrocytes within the cartilage template. Therefore, a perichondrial cell-specific mouse genetics tool would be essential to investigate the function of the perichondrium in skeletal development and regeneration.

In this study, we aimed to achieve two major goals. First, we aimed to define the cellular diversity of perichondrial cells, chondrocytes and their precursors of the pre-cartilaginous condensation and the cartilage template by single cell RNA-seq analyses. Second, by learning from novel marker genes and their expression patterns identified by single cell RNA-seq analyses, we sought to characterize previously unreported tamoxifen-inducible *creER* lines to identify potential cell-type specific inducible mouse genetics tools for fetal perichondrial cells and chondrocytes. We successfully characterized three new *creER* lines – *Hes1-creER*, *Dlx5-creER* and *Fgfr3-creER* – that might be instrumental for designing future studies. Our descriptive findings illustrate that perichondrial cells and chondrocytes collaborate in endochondral bone development, supporting the notion that these two types of cells are important contributors to the postnatal bone marrow stromal compartment.

## Results

### 1. Undifferentiated Hes1-creER^+^ mesenchymal cells provide precursors for skeletal progenitor cells within the cartilage template and the perichondrium of femurs

Endochondral bone development starts when undifferentiated mesenchymal cells make condensations, particularly around embryonic day (E) 10.5 in the mouse limb bud. These condensing mesenchymal cells express Sox9 and activate the transcription machinery for chondrocyte differentiation^17^. To define cellular heterogeneity of these undifferentiated mesenchymal cells in the condensation, we first undertook a single cell RNA-seq analysis. Mesenchymal cells contributing to the skeletal element of the limb are unanimously marked by a *cre* recombinase driven by a 2.4kb *Prrx1* promoter/enhancer^18^ (Fig.1a). We dissociated limb mesenchymal cells from *Prrx1-cre*; *R26R*^tdTomato^ mice at embryonic day 11.5 (E11.5), and isolated tdTomato^+^ cells by fluorescence-activated cell sorting (FACS) (Supplementary Fig.1a). We profiled 4,804 cells tdTomato^+^ cells using the 10X Chromium Single-Cell Gene Expression Solution platform. A graph-based clustering analysis using Seurat^19^ revealed 9 clusters, including two clusters of cells abundant in *Sox9* (Cluster 2,7, dotted contour) and six clusters of their surrounding cells (Cluster 0,1,3-6) (Fig.1b). The latter clusters were composed of mesenchymal cells expressing unique homeobox proteins, such as *Msx1* (Cluster 5,6), *Lhx9* (Cluster 5), *Irx3/5* (Cluster 3,7), *Meox2* (Cluster 0) and *Emx2* (Cluster 4) (Supplementary Fig.1b,c). In addition, *Shh* and *Fgf18* were expressed in cells in Cluster 6 and Cluster 0,1, respectively. Therefore, the limb bud skeletal element is constituted by distinct groups of mesenchymal cells potentially serving as precursors for chondrocytes and perichondrial cells.

**Figure 1.**
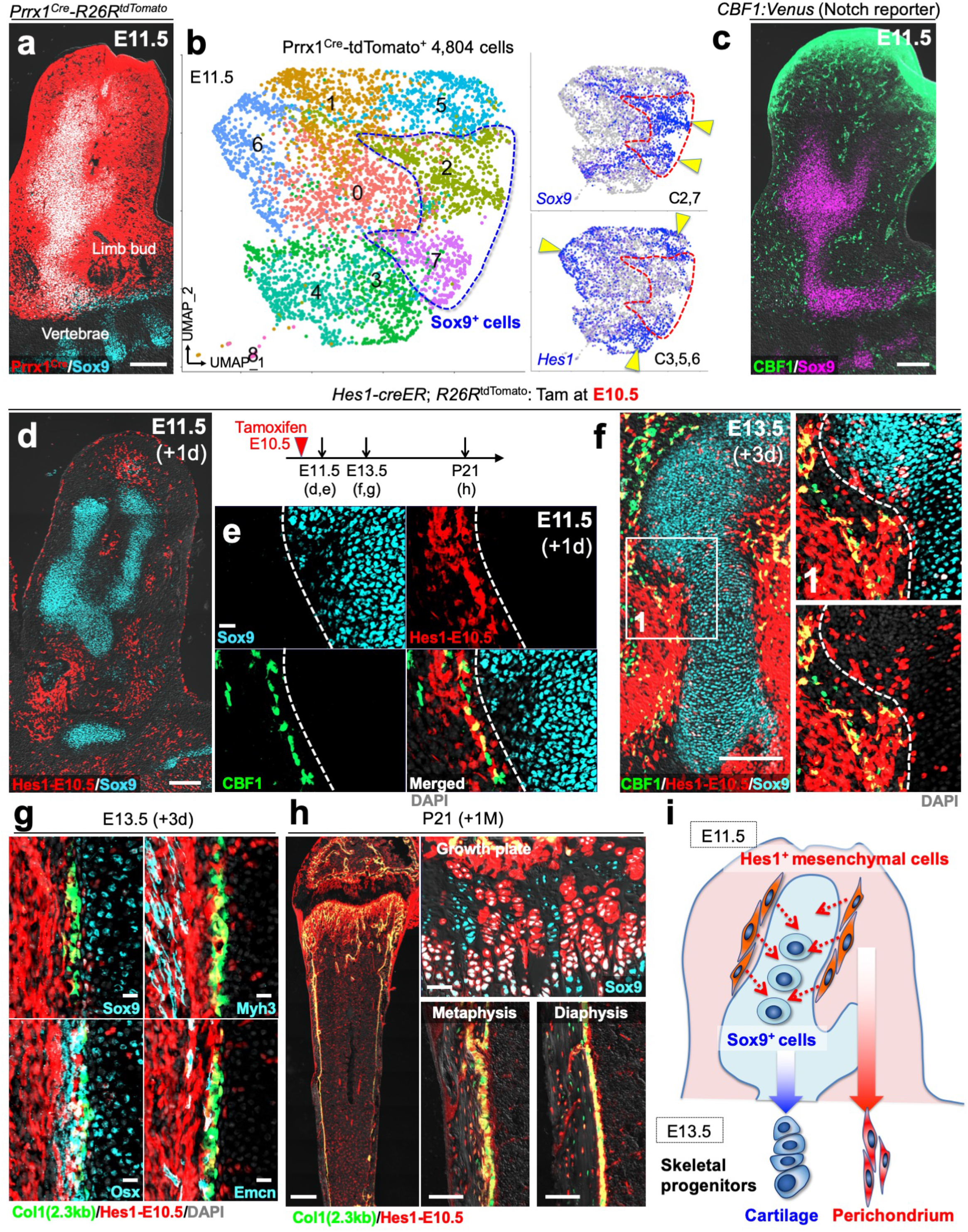
Undifferentiated Hes1^+^Sox9^neg^ mesenchymal cells provide precursors for skeletal progenitor cells within the cartilage template and the perichondrium. **(a,b)** Single cell RNA-seq analysis of *Prrx1-cre*-marked limb bud mesenchymal cells at E11.5. (a): *Prrx1-cre*; *R26R*^tdTomato^ femur stained for Sox9. Grey: DIC. Scale bar: 200µm. *n*=3 mice. (b): UMAP-based visualization of major classes of Prrx1^cre^-tdTomato^+^ cells (Cluster 0 – 8). Right panels: feature plots. Blue: high expression. Arrowheads: clusters in which a given gene is identified as a cell type-specific marker. Dotted contour: Sox9^+^ cells. *n*=4,804 cells. Pooled from *n*=5 mice. **(c)** Notch reporter *CBF1:H2B-Venus* femur immunostained for Sox9 at E11.5. Grey: DIC. Scale bar: 200µm. *n*=3 mice. **(d-h)** Cell-fate analysis of *Hes1-creER*^+^ mesenchymal cells of the condensation stage, pulsed at E10.5. *Hes1-creER*; *R26R*^tdTomato^ femurs carrying *CBF1:Venus* (e,f) or *Col1a1(2.3kb)-GFP* (g,h) reporters. (d): Limb bud at E11.5. Grey: DIC. Scale bar: 200µm. (e): Magnified view of mesenchymal condensation at E11.5. Grey: DAPI. Scale bar: 20µm. (f): Cartilage template at E13.5. Right panel: magnified view of Area (1). Dotted line: cartilage-perichondrium border. Grey: DIC. Scale bar: 200µm. (g): Cartilage-perichondrium immunostaining for Sox9, osterix (Osx), Myh3 and endomucin (Emcn). Grey: DAPI. Scale bar: 20µm. (h): Distal femur at P21 with growth plates on top. Whole bone (left), growth plates (upper right), endocortical marrow space at metaphysis (lower center) and diaphysis (lower right). Grey: DIC. Scale bar: 500µm (left panel), 50µm (upper right panel), 100µm (lower center and right panels). *n*=4 mice per each time point. **(i)** Diagram of Hes1^+^ mesenchymal cell fates. Hes1^+^Sox9^neg^ cells surrounding the condensation are *bona fide* skeletal progenitor cells during endochondral bone development, contributing to both chondrocytes and perichondrial cells, and subsequently to all limb skeletal cells.

In search for a cell-type specific marker for undifferentiated mesenchymal cells not yet committed to the chondrogenic pathway, we noticed that a Notch effector gene *Hes1* exhibited a pattern overlapping but negatively correlated with *Sox9*, identified as cell-type specific markers for Cluster 3,5,6 (Fig.1b, right panels). An RNAscope analysis revealed that *Hes1* was expressed in a manner surrounding the condensation expressing *Sox9* (Supplementary Fig.1d). To further validate Notch-responsive cells *in situ*, we took advantage of a Notch signaling reporter strain expressing a histone 2B (H2B)-bound Venus protein under a *C promoter binding factor 1 (CBF1)* promoter (*CBF1:H2B-Venus*)^20^. Interestingly, Notch-responsive Venus^bright^ cells were mostly located outside the SOX9^+^ domain of the condensation (Fig.1c). The discrepancy between *Hes1* expression pattern and CBF1:H2B-Venus reporter activities might indicate that Hes1^+^ cells might not be entirely Notch-responsive.

To study cell fates of these Hes1^+^ undifferentiated mesenchymal cells, we performed lineage-tracing experiments using a *Hes1-creER* knock-in allele^21^ that activates an *R26R*-tdTomato reporter in a tamoxifen-dependent manner. First, we pulsed *Hes1-creER*; *R26R*^tdTomato^ mice at E10.5 and analyzed these mice after 24 hours at E11.5, to define the characteristics of *Hes1-creER*^+^ cells (Hes1^CE^-E10.5 cells). Hes1^CE^-E10.5 cells were located throughout the limb bud except in SOX9^+^ pre-cartilaginous condensations, particularly in the area immediately adjacent to SOX9^+^ cells with abundant CBF1:H2B-Venus reporter activities (Fig.1d,e). Hes1^CE^-E10.5 cells expressed *Hes1*, but not *Sox9*, mRNAs at E11.5 (Supplementary Fig.2a). We also noticed that *Hes1-creER* simultaneously marked other types of cells outside the skeletal element of the limb bud, including MYH3^+^ skeletal muscle cells and EMCN^+^ endothelial cells (Supplementary Fig.2b). Taken together, *Hes1-creER* can mark mesenchymal cells in the SOX9-negative domain of the condensation upon tamoxifen injection.

Subsequently, we traced the fate of Hes1^+^ cells *in vivo* during endochondral bone development. After 3 days of chase at E13.5, Hes1^CE^-E10.5 cells contributed to both chondrocytes and perichondrial cells of the cartilage template; these cells became SOX9^+^ chondrocytes by translocating into the cartilage template, and a majority of perichondrial cells including those expressing CBF1:H2B-Venus (Fig.1f). In fact, Hes1^CE^-E10.5 cells contributed to all the layers of the perichondrium including OSX^+^ osteoblast precursors, in addition to MYH3^+^ skeletal muscle cells outside the perichondrium and EMCN^+^ endothelial cells (Fig.1g, Supplementary Fig.2c). After 9 days of chase at P0, Hes1^CE^-E10.5 cells contributed to a number of columnar chondrocytes of the growth plate, Col1a1(2.3kb)-GFP^+^ osteoblasts on the cortical and trabecular bone and stromal cells throughout the marrow space (Supplementary Fig.2d). Postnatally, Hes1^CE^-E10.5 cells contributed to the entire spectrum of the bone marrow stromal compartment; these cells became growth plate chondrocytes, osteoblasts both in the cortical and trabecular bone, and Cxcl12-GFP^high^ stromal cells throughout the marrow space both in the metaphysis and the diaphysis (Fig.1h, Supplementary Fig.2e,f). Skeletal muscle cells or endothelial cells that were initially marked by *Hes1-creER* at E10.5 did not contribute to these skeletal lineage cells, as cells marked by skeletal muscle-specific *cre* lines (*Acta1-cre*, *Myl1-cre* and *Mck-cre*) or an endothelial cell-specific *cre* line (*Tie2-cre*) did not contribute to chondrocytes, osteoblasts or bone marrow stromal cells at any time points (Supplementary Fig.3a,b).

Therefore, Hes1-creER^+^ mesenchymal cells of the condensation can behave as primordial skeletal progenitor cells, which robustly contribute to chondrocytes in the cartilage template, osteoblasts of the primary ossification center and the bone collar, and marrow stromal cells during endochondral bone development (Fig.1i).

### 2. *Hes1-creER* marks self-renewing osteo-stromal progenitor cells in the fetal femur

The perichondrium is defined when the fetal growth plate is established upon initial formation of hypertrophic chondrocytes^22^. Perichondrial cells are composed of multiple layers of mesenchymal cells with diverse functions; particularly, located on the innermost layer of the perichondrium adjacent to the prehypertrophic and hypertrophic layer is the osteogenic perichondrium, where cells become committed to the osteoblast lineage. First, we set out to define cellular heterogeneity of the fetal perichondrium by a single cell RNA-seq analysis. Both perichondrial cells and chondrocytes at the fetal stage are marked by a *cre* recombinase driven by a *Col2a1* promoter^12^ (Fig.2a). We dissociated limb mesenchymal cells of *Col2a1-cre*; *R26R*^tdTomato^ mice at E13.5, isolated tdTomato^+^ cells by FACS (Supplementary Fig.4a) and profiled 7,932 tdTomato^+^ cells using the 10X Chromium Single-Cell Gene Expression Solution platform. A graph-based clustering analysis revealed 11 clusters, including three clusters of chondrocytes abundant in *Sox9* (Cluster 0,1,3) and three clusters of perichondrial cells abundant in *Hes1* (Cluster 2,4,8) (Fig.2b). *Hes1* is most abundantly expressed in the fetal perichondrium^23^. In addition, Notch signaling is essential for maintaining skeletal progenitor cells by repressing Runx2 transcription activities^24^. Among the chondrocyte clusters, cells in Cluster 1 were enriched for *Col2a1*, *Fgfr3*, *Acan*, *Ihh* as well as *Runx2* and *Sp7*, representing relatively more mature chondrocytes (Fig.2b, Supplementary Fig.4b,c). Among the perichondrial cell clusters, cells in Cluster 4 were enriched for *Dlx5*, *Runx2*, *Prrx1* and *Sp7*, representing perichondrial cells partially committed to the osteoblast lineage (Fig.2b, Supplementary Fig.4b,c). Other perichondrial cells in Cluster 2 and 8 were enriched for *Igf1* and *Acta2*, respectively (Supplementary Fig.4b,c). Therefore, this analysis revealed the contiguous nature of the fetal chondrocyte-perichondrial cell lineage that shares an overlapping set of marker genes.

**Figure 2.**
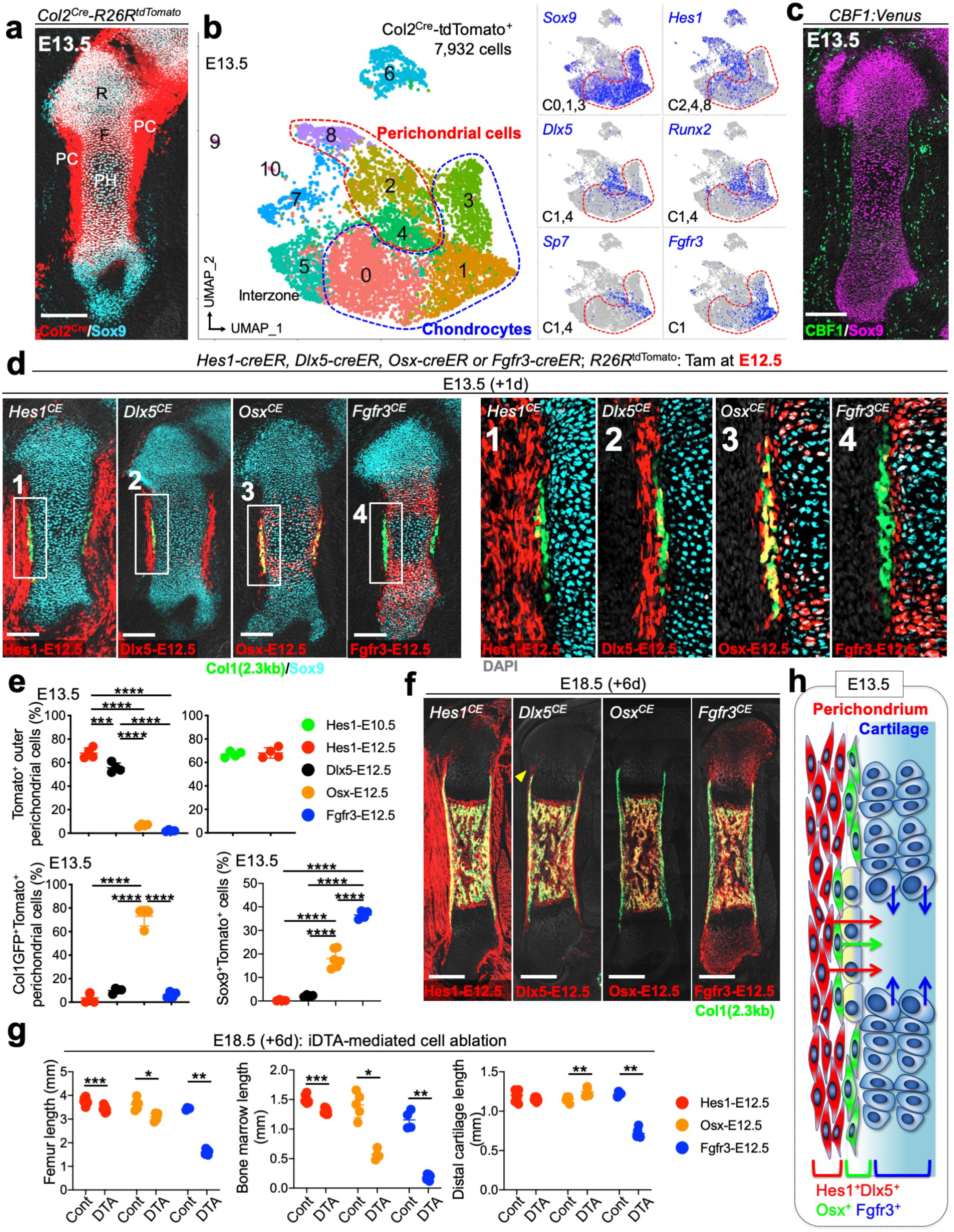
*Hes1-creER* marks self-renewing Dlx5^+^Osx^neg^ progenitor cells in the fetal perichondrium. **(a,b)** Single cell RNA-seq analysis of *Col2a1-cre*-marked chondrocytes and perichondrial cells at E13.5. (a): *Col2a1-cre*; *R26R*^tdTomato^ femur. R: Round layer. F: Flat layer. PH: Prehypertrophic layer. PC: Perichondrium. Grey: DIC. Scale bar: 200µm. *n*=3 mice. (b): UMAP-based visualization of major classes of Col2a1^cre^-tdTomato^+^ cells (Cluster 0 – 10). Red contour: perichondrial cells, Blue contour: chondrocytes. Right panels: feature plots of genes enriched in each cluster. Cluster 0,1,3: *Sox9*^+^, cluster 2,4,8: *Hes1*^+^, cluster 1: *Fgfr3*^+^. Blue: high expression. Dotted contour: chondrocytes. *n*=7,932 cells. Pooled from *n*=5 mice. **(c)** Notch reporter *CBF1:H2B-Venus* femur immunostained for Sox9 at E13.5. Grey: DIC. Scale bar: 200µm. *n*=3 mice. **(d-f)** Cell-fate analysis of *Hes1-creER*^+^, *Dlx5-creER*^+^, *Osx-creER*^+^ or *Fgfr3-creER*^+^ cells, pulsed at E12.5, carrying *Col1a1(2.3kb)-GFP* reporters. (d): Cartilage template at E13.5. Left panels: Grey: DIC. Scale bar: 200µm. Right panels: magnified view of (1-4). Grey: DAPI. *n*=4 mice per each group. (e): Upper: percentage of total tdTomato^+^ outer perichondrial cells among total outer perichondrial cells. *n*=4 mice per each group. \*\*\**p*<0.001, \*\*\*\**p*<0.0001, Hes1-E12.5 versus Fgfr3-E12.5, mean difference = 66.00, 95% confidence interval (59.56, 72.44); Hes1-E12.5 versus Osx-E12.5, mean difference = 61.29, 95% confidence interval (54.84, 67.73); Hes1-E12.5 versus Dlx5-E12.5, mean difference = 12.48, 95% confidence interval (6.034, 18.92); Fgfr3-E12.5 versus Osx-E12.5, mean difference = −4.711, 95% confidence interval (−11.15, 1.731); Fgfr3-E12.5 versus Dlx5-E12.5, mean difference = −53.52, 95% confidence interval (−59.96, − 47.08); Osx-E12.5 versus Dlx5-E12.5, mean difference = −48.81, 95% confidence interval (− 55.25, −42.37). Two-tailed, One-way ANOVA followed by Tukey’s post-hoc test (left). Two-tailed, Mann-Whitney’s *U*-test (right). Data are presented as mean ± s.d. Lower left: percentage of Col1a1(2.3kb)-GFP^+^tdTomato^+^ cells among total Col1a1(2.3kb)-GFP^+^ perichondrial osteoblasts. *n*=4 mice per each group. \*\*\*\**p*<0.0001, Hes1-E12.5 versus Fgfr3-E12.5, mean difference = −2.513, 95% confidence interval (−12.96, 7.933); Hes1-E12.5 versus Osx-E12.5, mean difference = −69.33, 95% confidence interval (−79.78, −58.88); Hes1-E12.5 versus Dlx5-E12.5, mean difference = −6.147, 95% confidence interval (−16.59, 4.299); Fgfr3-E12.5 versus Osx-E12.5, mean difference = −66.82, 95% confidence interval (−77.26, −56.37); Fgfr3-E12.5 versus Dlx5-E12.5, mean difference = −3.634, 95% confidence interval (−14.08, 6.813); Osx-E12.5 versus Dlx5-E12.5, mean difference = 63.18, 95% confidence interval (52.74, 73.63). Two-tailed, One-way ANOVA followed by Tukey’s post-hoc test. Data are presented as mean ± s.d. Lower right: Percentage of Sox9^+^tdTomato^+^ cells among total Sox9^+^ chondrocytes. *n*=4 (Hes1-E12.5, Dlx5-E12.5 and Fgfr3-E12.5), *n*=6 (Osx-E12.5). \*\*\*\**p*<0.0001, Hes1-E12.5 versus Fgfr3-E12.5, mean difference = −36.38, 95% confidence interval (−41.46, −31.30); Hes1-E12.5 versus Osx-E12.5, mean difference = −17.72, 95% confidence interval (−22.36, −13.08); Hes1-E12.5 versus Dlx5-E12.5, mean difference = −1.910, 95% confidence interval (−6.992, 3.171); Fgfr3-E12.5 versus Osx-E12.5, mean difference = 18.66, 95% confidence interval (14.02, 23.30); Fgfr3-E12.5 versus Dlx5-E12.5, mean difference = 34.47, 95% confidence interval (29.39, 39.55); Osx-E12.5 versus Dlx5-E12.5, mean difference = 15.81, 95% confidence interval (11.17, 20.45). Two-tailed, One-way ANOVA followed by Tukey’s post-hoc test. Data are presented as mean ± s.d. (f): Whole femurs at E18.5. Arrowhead: groove of Ranvier. Grey: DIC. Scale bar: 500µm. *n*=4 mice per each group. **(g)** Inducible partial cell ablation of *Hes1-creER*^+^, *Osx-creER*^+^ or *Fgfr3-creER*^+^ cells using an inducible Diphtheria toxin fragment A (iDTA) allele. Quantification of femur length, bone marrow length and distal cartilage length. *n*=7 (Hes1-Cont), 6 (Hes1-DTA, Fgfr3-DTA), 5 (Osx-Cont), 4 (Osx-DTA, Fgfr3-Cont). \**p*<0.05, \*\**p*<0.01, \*\*\**p*<0.001, two-tailed, Mann-Whitney’s *U*-test. Data are presented as mean ± s.d. **(h)** Diagram of Hes1-E12.5, Dlx5-E12.5, Osx-E12.5, or Fgfr3-E12.5, cell fates. *Hes1-creER* and *Dlx5-creER*, but not *Osx-creER* or *Fgfr3-creER*, can mark a self-renewing population of fetal perichondrial cells with robust capability to contribute to cortical and trabecular bone compartments.

We further constructed single-cell trajectories of tdTomato^+^ cells using Monocle 2^25,26^. This unsupervised analysis predicted a branched trajectory starting from *Prrx1*^+^ cells and ending either at *Ihh*^+^ pre-hypertrophic chondrocytes or *Acta2*^+^ perichondrial cells, indicating that chondrocytes and perichondrial cells represent discrete cell fates of Prrx1^+^ cells in the primordium (Supplementary Fig.4d). We subsequently analyzed the branch point using branched expression analysis modeling (BEAM) and heatmap-based visualization of representative branch-dependent genes, to define genes involved with two different cell fate choices (Supplementary Fig.4d). Hierarchical clustering revealed that a chondrocyte fate and a perichondrial cell fate represent distinct transcriptomic paths; a chondrocyte fate was associated with sequential upregulation of *Bmp2*^27^, *Sox9*^28^ and *Runx2*^29^, whereas a perichondrial cell fate was associated with upregulation of a Wnt-target gene, *Axin2*^30^ (Supplementary Fig.4e). Therefore, these trajectory analyses support the notion that perichondrial cells represent a discrete state emanating as a cell fate alternative to chondrocytes.

We noticed that the expression pattern of *Hes1* was negatively correlated with that of *Fgfr3*. To reveal the relationship between *Hes1* and *Fgfr3*, we performed a MAGIC imputation analysis^31^. As expected, cells expressing *Fgfr3* expressed *Sox9* at a unanimously high level (Supplementary Fig.4f). In contrast, *Hes1* expression demonstrated an overall negative correlation with *Fgfr3* expression. In fact, an RNAscope analysis confirmed that *Hes1* was expressed in the perichondrium in a manner distinct from *Fgfr3*, which was predominantly expressed within the cartilage template (Supplementary Fig.5a,b).

Further analysis with *CBF1:H2B-Venus* mice revealed that Notch-responsive Venus^bright^ cells were located outside SOX9^+^ cells in a pattern surrounding the fetal cartilage at E13.5 (Fig.2c). We also validated in situ expression patterns of *Fgfr3* by taking advantage of an *Fgfr3-GFP* bacterial artificial chromosome (BAC) transgenic strain. Fgfr3-GFP^+^ cells were found within the SOX9^+^ domain of the fetal cartilage template, in a pattern distinct from CBF1:H2B-Venus^bright^ cells (Supplementary Fig.5b). Fgfr3-GFP^+^ cells expressed *Fgfr3* mRNA (Supplementary Fig.5b). Therefore, Notch effector gene *Hes1* is expressed in a pattern largely distinct from that of *Fgfr3*.

Subsequently, to delineate cell fates of perichondrial cell populations, we took advantage of tamoxifen-inducible *creER* lines – *Hes1-creER, Dlx5-creER*^32^ and *Osx-creER*^6^ – whose expression is regulated by cell-type specific marker genes identified by the above-mentioned single cell RNA-seq analysis. We also took advantage of an *Fgfr3-creER* P1-derived artificial chromosome (PAC) transgenic line^33^ to contrast their cell fates with those of cells within the cartilage template. First, we pulsed these mice carrying *R26R^tdTomato^* and *Col1a1(2.3kb)-GFP* at E12.5, and analyzed these mice after 24 hours at E13.5, to define the characteristics of *Hes1-creER^+^*, *Dlx5-creER^+^*, *Osx-creER^+^* and *Fgfr3-creER^+^* cells at this stage (Hes1^CE^-E12.5, Dlx5^CE^-E12.5, Osx^CE^-E12.5 and Fgfr3^CE^-E12.5 cells, respectively). Hes1^CE^-E12.5 cells were located outside the SOX9^+^ domain of the cartilage template with minimal overlap with Col1a1(2.3kb)-GFP^+^ osteoblasts in the perichondrium (Fig.2d). Some of these cells also expressed CBF1:H2B-Venus (Supplementary Fig.5a). Hes1^CE^-E12.5 cells were also found outside the skeletal element among MYH3^+^ skeletal muscle cells and EMCN^+^ endothelial cells (Supplementary Fig.5c). Dlx5^CE^-E12.5 cells occupied the domain of the perichondrium equivalent to Hes1^CE^-E12.5 cells, but in a more specific pattern without any overlap with MYH3^+^ skeletal muscle cells or EMCN^+^ endothelial cells (Fig.2d, Supplementary Fig.5d). In contrast, Osx^CE^-E12.5 cells occupied the perichondrium in a pattern distinct from Dlx5^CE^-E12.5 cells, with the majority of these cells either overlapping or adjoining with Col1a1(2.3kb)-GFP^+^ osteoblasts (Fig.2d). A substantial number of Osx^CE^-E12.5 cells were also located within the cartilage as reported previously^6^. Fgfr3^CE^-E12.5 cells were predominantly located within the cartilage with minimal overlap with Col1a1(2.3kb)-GFP^+^ perichondrial osteoblasts; a small number of Fgfr3^CE^-E12.5 cells were also found in the perichondrium (Fig.2d).

Quantification of tdTomato^+^ perichondrial cells in each model revealed that *Hes1-creER* and *Dlx5-creER* could mark undifferentiated precursor populations of perichondrial cells. The number of Hes1^CE^-E12.5 and Dlx5^CE^-E12.5 cells in the perichondrium was comparable at E13.5 (Fig.2e, top left panel). In addition, Hes1^CE^-E12.5 and Dlx5^CE^-E12.5 perichondrial cells were distinct from perichondrial osteoblasts, as virtually no fraction of these cells was Col1a1(2.3kb)-GFP^+^ (Fig.2e, bottom left panel). Importantly, Hes1^+^ descendants continued to express *Hes1* in the perichondrium, as the relative fraction of Hes1^CE^-E10.5 and Hes1^CE^-E12.5 cells in the perichondrium remained comparable at E13.5 (Fig.2e, top right panel). Further quantification of tdTomato^+^SOX9^+^ cells revealed that Hes1^CE^-E12.5 and Dlx5^CE^-E12.5 cells were distinct from chondrocytes (Fig.2e, bottom right panel). Therefore, *Hes1-creER* and *Dlx5-creER* can mark undifferentiated perichondrial cells that are largely distinct from chondrocytes and perichondrial cells marked by *Osx-creER* or *Fgfr3-creER* at this stage.

CBF1 (encoded by *Rbpj*) and its transcriptional target *Hes1* regulate differentiation and hypertrophy of chondrocytes^34,35^. Analysis of CBF1:H2B-Venus mice at a later stage of E15.5 and P5 revealed that Venus^bright^ cells were also present in proliferating and articular chondrocytes; however, *Hes1-creER* did not mark any chondrocyte within the growth plate at E15.5 and P5 upon a pulse at E14.5 and P3, respectively, (Supplementary Fig.5e,f), revealing the dissociation between CBF1:H2B-Venus and *Hes1-creER* activities at a later stage.

Subsequently, we traced the fate of these distinct populations of perichondrial cells during formation of the primary ossification center and the marrow space. After 3 days of chase at E15.5, Hes1^CE^-E12.5 cells, Dlx5^CE^-E12.5 cells and Osx^CE^-E12.5 cells contributed to mesenchymal cells within the primary ossification center as well as Col1a1(2.3kb)-GFP^+^ osteoblasts in the bone collar (Supplementary Fig.6a). Fgfr3^CE^-E12.5 cells expanded within the cartilage template as SOX9^+^ chondrocytes, while exuberantly translocating into the nascent primary ossification center (Supplementary Fig.6a). After 6 days of chase at E18.5, these cells – Hes1^CE^-E12.5, Dlx5^CE^-E12.5, Osx^CE^-E12.5 and Fgfr3^CE^-E12.5 cells – contributed to Col1a1(2.3kb)-GFP^+^ osteoblasts in the trabecular bone (Fig.2f, Supplementary Fig.6b) and Cxcl12-GFP^high^ marrow stromal cells in the marrow space (Supplementary Fig.6c). Importantly, unlike those pulsed at an earlier stage, Hes1^CE^-E12.5 cells did not contribute to SOX9^+^ chondrocytes in the growth plate (Fig.2f, Supplementary Fig.6b). Therefore, mesenchymal cells within the primary ossification center might have dual origins in the perichondrium and the cartilage template.

Additionally, we noticed a notable difference with regard to how these cells contributed to formation of the perichondrium, the periosteum and the bone collar. Fgfr3^CE^-E12.5 cells did not contribute to the perichondrium, the bone collar or the cortical bone at the subsequent stages (Fig.2f, Supplementary Fig.6b). Osx^CE^-E12.5 cells did not remain in the perichondrium as reported previously^5^, therefore not robustly contributing to the bone collar or the cortical bone at the subsequent stages (Fig.2f, Supplementary Fig.6b). In contrast, Dlx5^CE^-E12.5 cells maintained themselves within the perichondrium including the groove of Ranvier (Fig.2f, Supplementary Fig.6b, arrowheads), and robustly contributed to the bone collar and the cortical bone. Hes1^CE^-E12.5 cells demonstrated a pattern similar to Dlx5^CE^-E12.5 cells without notable contribution to growth plate chondrocytes (Fig.2f, Supplementary Fig.6b). Therefore, *Hes1-creER* and *Dlx5-creER*, but not *Osx-creER* or *Fgfr3-creER*, can mark a self-renewing population of fetal perichondrial cells with robust capability to contribute to cortical and trabecular bone compartments. Although whether these cell populations are fully overlapping cannot be formally demonstrated, it is important to note that Hes1-creER^+^ and Dlx5-creER^+^ cells behave in a similar manner.

To clarify the uniqueness of the *Fgfr3-creER* line that predominantly marks chondrocyte-like cells within the cartilage template, we also analyzed a well-characterized *Col2a1-creER* line^13^. Analysis of *Col2a1-creER; R26R^tdTomato^* mice at E13.5 after 24 hours of pulse at E12.5 revealed that Col2a1^CE^-E12.5 cells were present not only in the cartilage template but also in the perichondrium (Supplementary Fig.6d). After 2 weeks of chase at P7, Col2a1^CE^-E12.5 cells contributed to essentially all perichondrial cells and growth plate chondrocytes, whereas Fgfr3^CE^-E12.5 cells contributed to virtually no perichondrial cells and to a certain fraction of growth plate chondrocytes (Supplementary Fig.6d). Therefore, *Fgfr3-creER* marks a specific subset of cells that are marked by *Col2a1-creER*.

We further set out to define the functional significance of these cell populations in endochondral bone development. For this purpose, we performed inducible cell ablation experiments using a *Rosa26-iDTA* (inducible diphtheria toxin fragment A) allele, which was crossed with aforementioned *creER* lines. This iDTA-mediated approach can only partially ablate target cells of interest upon tamoxifen injection^36,37^. We pulsed *Hes1-creER*, *Osx-creER or Fgfr3-creER*; *Rosa26*^lsl-tdTomato/+^ (Cont) and *Hes1-creER*, *Osx-creER or Fgfr3-creER*; *Rosa26*^lsl-tdTomato/iDTA^ (DTA) mice at E12.5, and analyzed these mice at 6 days later at E18.5. All of these cells – Hes1^CE^-E12.5, Osx^CE^-E12.5 and Fgfr3^CE^-E12.5 cells – were functionally significant contributors to bone development, as the femur length was significantly reduced in all groups owing to iDTA-mediated cell ablation (Fig.2g, left panel, Supplementary Fig.6e). Notably, ablation of Hes1^+^ cells selectively impaired the formation of bone marrow without affecting the cartilage, as the bone marrow length, but not the cartilage length, was significantly reduced in the Hes1^CE^-E12.5 DTA group (Fig.2g, middle and right panel).

Therefore, all of these cells – Hes1-creER^+^, Dlx5-creER^+^, Osx-creER^+^ and Fgfr3-creER^+^ cells – provide the functionally significant pathway to generate skeletal lineage cells within the nascent marrow space (Fig.2h).

### 3. Hes1-creER^+^ and Dlx5-creER^+^ perichondrial cells substantially contribute to the postnatal bone marrow stromal compartment of femurs

We further set out to define how Hes1-creER^+^ and Dlx5-creER^+^ cells contribute to the postnatal bone marrow stromal compartment. For this purpose, we chased Hes1^CE^-E12.5, Dlx5^CE^-E12.5, Osx^CE^-E12.5 and Fgfr3^CE^-E12.5 until postnatal day (P) 21. Hes1^CE^-E12.5 and Dlx5^CE^-E12.5 cells predominantly contributed to the diaphyseal bone marrow stromal compartment; these cells became Col1a1(2.3kb)-GFP^+^ osteoblasts both in the cortical and trabecular bone and Cxcl12-GFP^high^ stromal cells (Fig.3a,b,e,f, Supplementary Fig.7a,b). In contrast, Osx^CE^-E12.5 cells contributed to osteoblasts and marrow stromal cells in a more limited domain of the diaphyseal bone marrow, particularly on the proximal end of the marrow space (Fig.3c,g, Supplementary Fig.7c, ref.^5,7,38^). Fgfr3^CE^-E12.5 cells became cortical and trabecular osteoblasts as well as Cxcl12-GFP^high^ stromal cells predominantly in the metaphyseal marrow space, in a mutually exclusive manner to aforementioned cell types (Fig.3d,h, Supplementary Fig.7d). Interestingly, Hes1^CE^-E14.5 cells contributed to only a small fraction of osteoblasts or Cxcl12-GFP^high^ stromal cells; instead, *Hes1-creER* marked cells of the endothelial lineage in the nascent marrow space and contributed to a large number of sinusoidal endothelial cells in the postnatal marrow space (Supplementary Fig.8).

**Figure 3.**
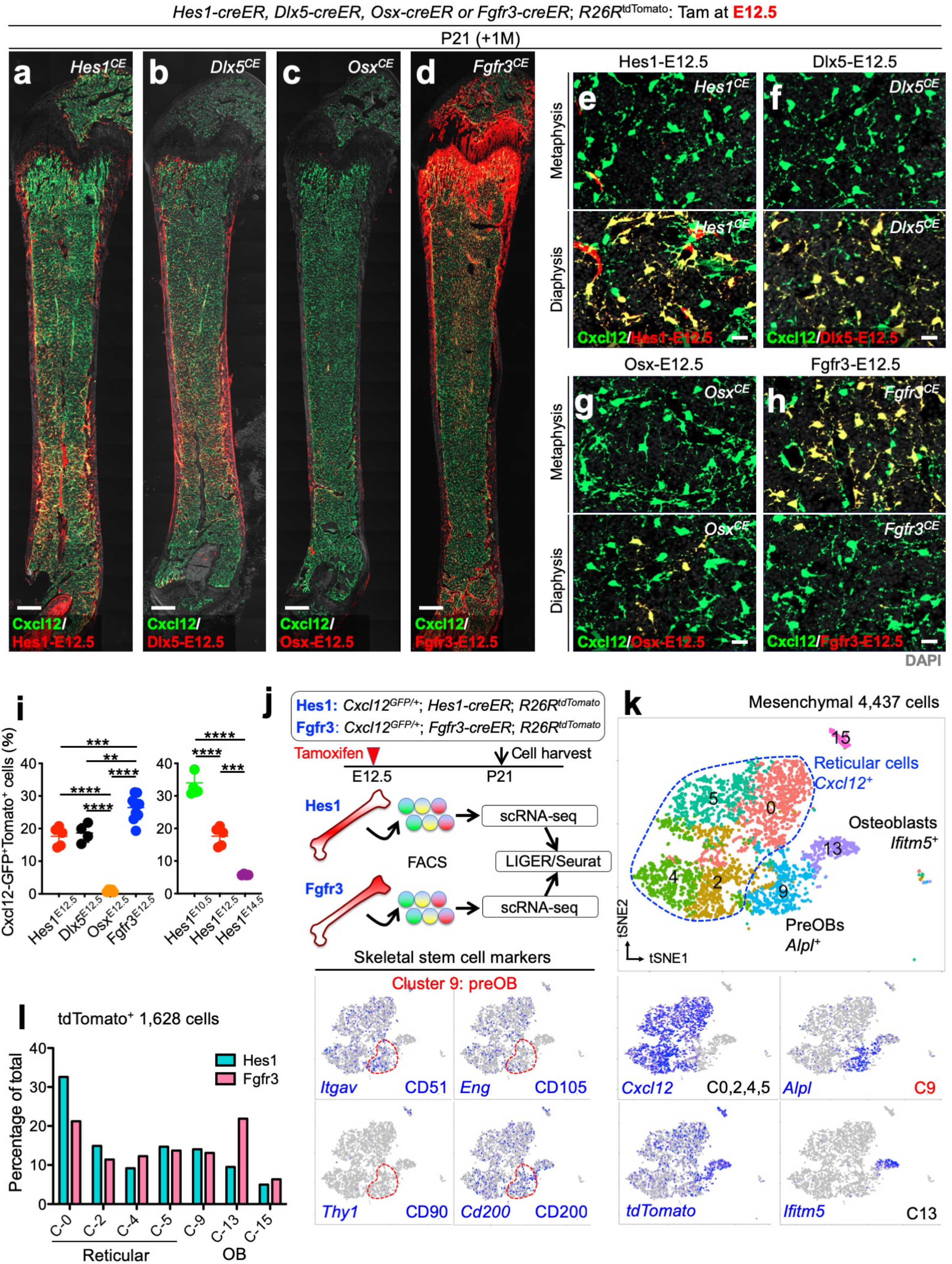
Hes1^+^Dlx5^+^Osx^neg^ perichondrial cells substantially contribute to the postnatal bone marrow stromal compartment. **(a-i)** Contribution of fetal *Hes1-creER*^+^ cells, *Dlx5-creER*^+^ cells, *Osx-creER*^+^ cells or *Fgfr3-creER*^+^ cells (pulsed at E12.5) to Cxcl12-GFP^+^ bone marrow stromal cells at P21. *Cxcl12^GFP/+^*; *Hes1-creER*; *R26R^tdTomato^* (a,e), *Cxcl12^GFP/+^*; *Dlx5-creER*; *R26R^tdTomato^* (b,f), *Cxcl12^GFP/+^*; *Osx-creER*; *R26R^tdTomato^* (c,g) or *Cxcl12^GFP/+^*; *Fgfr3-creER*; *R26R^tdTomato^* (d,h) femurs with growth plates on top. (a-d): Whole bone, upper panels in (e-h): metaphyseal bone marrow, lower panels in (e-h): diaphyseal bone marrow. Grey: DIC (a-d), DAPI (e-h). Scale bar: 500µm (a-d), 20µm (e-h). *n*=4 mice per group. (i): Percentage of Cxcl12-GFP^high^tdTomato^+^ cells per total Cxcl12-GFP^high^ cells. *n*=5 (Hes1-E12.5), *n*=4 (Dlx5-E12.5), *n*=5 (Osx-E12.5), *n*=10 (Fgfr3-E12.5), *n*=6 (Hes1-E10.5), *n*=4 (Hes1-E14.5) mice per group. Left: \*\**p*<0.01, \*\*\**p*<0.001, \*\*\*\**p*<0.0001, Hes1-E12.5 versus Dlx5-E12.5, mean difference = −1.050, 95% confidence interval (−6.686, 4.586); Hes1-E12.5 versus Osx-E12.5, mean difference = 16.65, 95% confidence interval (11.34, 21.96); Hes1-E12.5 versus Fgfr3-E12.5, mean difference = −8.840, 95% confidence interval (− 13.44, −4.238); Dlx5-E12.5 versus Osx-E12.5, mean difference = 17.70, 95% confidence interval (12.06, 23.34); Dlx5-E12.5 versus Fgfr3-E12.5, mean difference = −7.790, 95% confidence interval (−12.76, −2.819); Osx-E12.5 versus Fgfr3-E12.5, mean difference = −25.49, 95% confidence interval (−30.09, −20.89). Two-tailed, One-way ANOVA followed by Tukey’s post-hoc test. Data are presented as mean ± s.d. Right: \*\*\**p*<0.001, \*\*\*\**p*<0.0001, Hes1-E10.5 versus Hes1-E12.5, mean difference = 16.42, 95% confidence interval (11.40, 21.44); Hes1-E10.5 versus Hes1-E14.5, mean difference = 28.23, 95% confidence interval (22.88, 33.58); Hes1-E12.5 versus Hes1-E14.5, mean difference = 11.82, 95% confidence interval (6.258, 17.38). Two-tailed, One-way ANOVA followed by Tukey’s post-hoc test. Data are presented as mean ± s.d. **(j-l)** Combined lineage-tracing and single cell RNA-seq analyses of Cxcl12-GFP^+^ bone marrow stromal cells and tdTomato^+^ mesenchymal cells derived from fetal Hes1^+^ perichondrial cells and Fgfr3^+^ chondrocytes. (j): Both single and double positive cells (GFP^high^, tdTomato^+^ and GFP^high^tdTomato^+^) were FACS-isolated from *Cxcl12^GFP/+^*; *Hes1-creER*; *R26R*^tdTomato^ and *Cxcl12^GFP/+^*; *Fgfr3-creER*; *R26R*^tdTomato^ femurs at P21, pulsed at E12.5 (Hes1-E12.5 or Fgfr3-E12.5). Two single cell datasets were integrated by LIGER. (k): *t*-SNE-based visualization of major classes of mesenchymal Cxcl12-GFP^high^ and/or tdTomato^+^ (Hes1-E12.5 or Fgfr3-E12.5) cells (Cluster 0, 2, 4, 5, 9, 13, 15). Lower panels: feature plots. Blue: high expression. Blue contour: *Cxcl12*^+^ reticular cells. *n*=4,437 cells. Pooled from *n*=4 mice per group. (l): Cells colored by conditions (Hes1-E12.5, Fgfr3-E12.5), and percentage of cells in each cluster among total cells, demonstrated for each condition. *n*=1,628 cells (tdTomato^+^).

To assess quantitatively how perichondrial cells and chondrocytes contribute differentially to Cxcl12-GFP^high^ stromal cells at P21, we performed flow cytometry analysis of cells isolated from femurs of triple transgenic mice carrying a Cxcl12-GFP reporter (Fig.3i, Supplementary Fig.9a). First, undifferentiated Hes1-creER^+^ mesenchymal cells in the condensation demonstrated a robust potential as stromal progenitor cells, as Hes1^CE^-E10.5 cells contributed to a substantial fraction of Cxcl12-GFP^high^ stromal cells (34.0±4.0%). Second, *Hes1-creER* and *Dlx5-creER* could mark comparable perichondrial progenitor cell populations at E12.5, as Hes1^CE^-E12.5 and Dlx5^CE^-E12.5 cells contributed to an equivalent fraction of Cxcl12-GFP^high^ stromal cells (17.6±2.9% and 18.6±4.6%, respectively). Third, Hes1-creER^+^ and Dlx5-creER^+^ cells, but not Osx-creER^+^ cells, represent a substantial source of bone marrow stromal cells, as Osx^CE^-E12.5 cells contributed to only a small fraction of Cxcl12-GFP^high^ stromal cells (1.0±0.4%). Fourth, fetal Fgfr3-creER^+^ cells may provide a primary source of bone marrow stromal cells and osteoblasts, as Fgfr3^CE^-E12.5 cells contributed to a higher fraction of Cxcl12-GFP^high^ stromal cells (26.4±3.7%) than Hes1^CE^-E12.5 or Dlx5^CE^-E12.5 cells did. Additional analysis of transgenic mice carrying a Col1(2.3kb)-GFP reporter revealed that Fgfr3^CE^-E12.5 cells contributed to a significantly higher fraction of Col1(2.3kb)-GFP^+^ cells (19.6±3.0%) than Hes1^CE^-E12.5 cells did (5.5±1.7%) (Supplementary Fig.9b). Lastly, Hes1-creER^+^ cells may contribute to bone marrow stromal cells in a time-specific fashion, as Hes1^CE^-E14.5 cells contributed to a much less fraction of Cxcl12-GFP^high^ stromal cells than Hes1^CE^-E12.5 cells did (5.8±0.2%).

To interrogate the characteristics of Hes1-creER^+^ cell-derived bone marrow stromal cells, we further performed an integrative single cell RNA-seq analysis combining two datasets (Cxcl12-GFP^+^||Hes1^CE^E12.5-tdTomato^+^ and Cxcl12-GFP^+^||Fgfr3^CE^E12.5-tdTomato^+^) at P21 using LIGER (Fig.3j). Mesenchymal GFP^+^||tdTomato^+^ cells clustered into several groups, including four groups of *Cxcl12*^+^ reticular cells (Cluster 0,2,4,5), *Alpl*^+^ pre-osteoblasts (Cluster 9) and *Ifitm5*^+^ mature osteoblasts (Cluster 13) (Fig.3k, Supplementary Fig.10a-c). Interestingly, cells in Cluster 9 also expressed *Itgav* (encoding CD51) and *Cd200* (CD200), but not *Thy1* (CD90) or *Eng* (CD105), indicating that these cells might coincide with previously reported mouse skeletal stem cells (mSSCs)^39^. We further subclustered tdTomato^+^ cells for the following analysis. Hes1^CE^-E12.5 and Fgfr3^CE^-E12.5 cells were distributed across all seven clusters including Cluster 9 that was enriched for mSSC-like cells. Flow cytometry analysis revealed that a majority of both Cxcl12-GFP^+^Hes1^CE^-E12.5 and Cxcl12-GFP^+^Fgfr3^CE^-E12.5 cells expressed CD200, but not CD90 (Supplementary Fig.10e). We noticed that Fgfr3^CE^-E12.5 cells were relatively overrepresented in a mature osteoblast cluster (Cluster 13, *Ifitm5*^+^*Bglap*^+^) (Fig.3l), suggesting that Fgfr3^+^ chondrocyte-like cells might preferentially contribute to mature osteoblasts within the stromal compartment. Furthermore, Fgfr3^CE^-E12.5 cells exhibited higher expression of chondrocyte signature genes, including *Col2a1*, *Col10a1*, *Acan*, *Matn3* as well as osteoblast signature genes, including *Prrx1* and *Sp7*, both in terms of the percentage of cells expressing given genes and the expression level, compared with Hes1^CE^-E12.5 cells (Supplementary Fig.10d). In contrast, Hes1^CE^-E12.5 cells exhibited higher expression of adipocyte signature genes including *Fabp4* and *Cebpb* (Supplementary Fig.10d). Therefore, BMSCs originating from two different sources of the perichondrium and the cartilage template might maintain at least some of the unique gene expression signatures after translocating into the stromal compartment.

Taken together, these findings reveal that Hes1-creER^+^, Dlx5-creER^+^ and Fgfr3-creER^+^ cells, but not Osx-creER^+^ cells, in the fetal stage contribute to a substantial fraction of the bone marrow stromal compartment during the postnatal stage.

### 4. Hes1-creER^+^ perichondrial cell-derived marrow stromal cells persist in the diaphysis and participate locally in osteogenesis of femurs

Lastly, we set out to determine the functional significance of Hes1-creER^+^ cell-derived cells in the bone marrow stromal compartment. To this end, we first performed CFU-F assays of Hes1^CE^-E12.5, Fgfr3^CE^-E12.5 and Osx ^CE^-E12.5 cells. At an early postnatal stage of P7, Hes1^CE^-E12.5 cells contributed to a higher faction of CFU-Fs than Fgfr3^CE^-E12.5 cells did; importantly, Osx^CE^-E12.5 cells contributed to no CFU-F (Hes1^CE^-E12.5: 33.8±7.2%, Fgfr3^CE^-E12.5: 13.2±7.0%, Osx^CE^-E12.5: 0.0±0.0%, Fig.4a,c). By contrast, Fgfr3^CE^-E12.5 cells contributed to a higher faction of CFU-Fs than Hes1^CE^-E12.5 cells did at P21 (Hes1^CE^-E12.5: 13.7±7.2%, Fgfr3^CE^-E12.5: 57.1±11.3%, Fig.4b,c). This inversed relationship was associated with a significant decrease of Hes1^CE^-E12.5 CFU-Fs and a significant increase of Fgfr3^CE^-E12.5 CFU-Fs during postnatal bone growth (Fig.4c). Isolation of individual tdTomato^+^ colonies and their subsequent passaging revealed that a small fraction of both Hes1^CE^-E12.5 and Fgfr3^CE^-E12.5 CFU-Fs could survive at least for ten generations (Fig.4d). Therefore, Hes1-creER^+^ cell-derived cells make a substantial contribution to a colony-forming fraction of BMSCs particularly during the early postnatal stage. Interestingly, BMSCs derived from Fgfr3-creER^+^ cells make a progressive contribution to a colony-forming fraction, and compose a majority of these clonogenic cells in the later postnatal stage.

**Figure 4.**
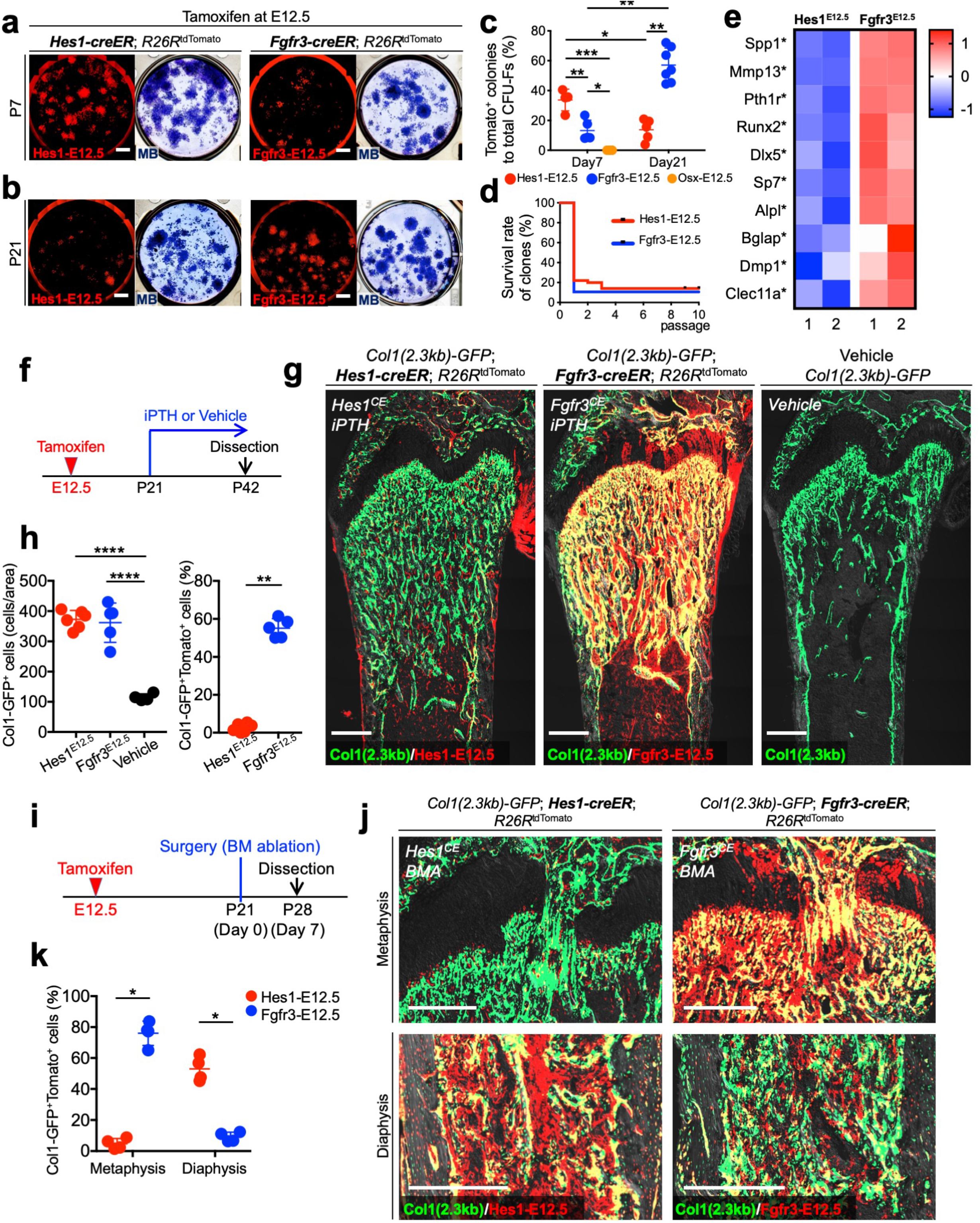
Site-specific roles of fetal perichondrium-derived and chondrocyte-derived mesenchymal stromal cells in osteoblast differentiation. **(a-d)** Colony-forming unit fibroblast (CFU-F) assay of E12.5-pulsed *Hes1-creER*; *R26R*^tdTomato^ (left panels), *Fgfr3-creER*; *R26R*^tdTomato^ (right panels) and *Osx-creER*; *R26R*^tdTomato^ bone marrow cells, at P7 (a) and P21 (b). MB: Methylene blue staining. Scale bar: 5mm. (c): Percentage of tdTomato^+^ colonies among total CFU-Fs. Hes1-E12.5: *n*=4 (P7), *n*=5 (P21), Fgfr3-E12.5: *n*=5 (P7), *n*=7 (P21), Osx-E12.5: *n*=3 (P7) mice. Hes1-E12.5 versus Fgfr3-E12.5 versus Osx-E12.5 (Day7): \**p*<0.05, \*\**p*<0.01, \*\*\**p*<0.001, Hes1-E12.5 versus Fgfr3-E12.5, mean difference = 20.51, 95% confidence interval (8.789, 32.23); Hes1-E12.5 versus Osx-E12.5, mean difference = 33.75, 95% confidence interval (20.40, 47.10); Fgfr3-E12.5 versus Osx-E12.5, mean difference = 13.24, 95% confidence interval (0.4796, 26.00). Two-tailed, One-way ANOVA followed by Tukey’s post-hoc test. Data are presented as mean ± s.d. Hes1-E12.5 versus Fgfr3-E12.5 versus Osx-E12.5 (Day7), Hes1-E12.5 (Day7 versus Day21), Fgfr3-E12.5 (Day7 versus Day21): \**p*<0.05, \*\**p*<0.01, two-tailed, Mann-Whitney’s *U*-test. Data are presented as mean ± s.d. (d): Survival curve of Hes1-E12.5 and Fgfr3-E12.5 individual tdTomato^+^ clones (P21) over 10 passages. *n*=50 (Hes1-E12.5), *n*=60 (Fgfr3-E12.5) clones per group. Gehan-Breslow-Wilcoxon test. **(e)** Comparative bulk RNA-seq analysis of Cxcl12-GFP^high^ stromal cells derived from fetal Hes1^+^ perichondrial cells and Fgfr3 ^+^ chondrocytes. Heatmap of representative differentially expressed genes (DEGs) associated with osteoblast differentiation. Star: statistically significant difference between Hes1-E12.5 and Fgfr3-E12.5. *n*=2 biological replicates (Hes1-E12.5: pooled from *n*=4 mice, Fgfr3-E12.5: pooled from *n*=5 mice). **(f-h)** Differential responses of mesenchymal stromal cells derived from Hes1^+^ perichondrial cells and Fgfr3^+^ chondrocytes to intermittent administration of parathyroid hormone (PTH). (g): *Col1a1(2.3kb)-GFP*; *Hes1-creER*; *R26R*^tdTomato^ (Hes1-E12.5, left), *Col1a1(2.3kb)-GFP*; *Fgfr3-creER*; *R26R*^tdTomato^ (Fgfr3-E12.5, center), and *Col1a1(2.3kb)-GFP*; *R26R*^tdTomato^ (Control, right) distal femurs, with iPTH or Vehicle for 3 weeks from P21 to P42. Grey: DIC. Scale bar: 500µm. (h): Quantification of Col1a1(2.3kb)-GFP^+^ cells (left) and Col1a1(2.3kb)-GFP^+^tdTomato^+^ cells (right) in the metaphysis. *n*=6 (Hes1-E12.5), *n*=5 (Fgfr3-E12.5), *n*=4 (Control) mice. \*\*\*\**p*<0.0001, Hes1-E12.5 versus Fgfr3-E12.5, mean difference = 10.5, 95% confidence interval (−58.2, 79.2); Hes1-E12.5 versus Vehicle, mean difference = 257.8, 95% confidence interval (184.5, 331); Fgfr3-E12.5 versus Vehicle, mean difference = 247.3, 95% confidence interval (171.1, 323.4). Two-tailed, One-way ANOVA followed by Tukey’s post-hoc test (left). \*\**p*<0.01, two-tailed, Mann-Whitney’s *U*-test (right). Data are presented as mean ± s.d. **(i-k)** Injury-responsive osteogenesis of mesenchymal stromal cells derived from fetal perichondrial cells (Hes1-E12.5) and chondrocytes (Fgfr3-E12.5) cells. (j): Bone marrow ablation for trabecular bone regeneration in *Col1a1(2.3kb)-GFP*; *Hes1-creER; R26R*^tdTomato^ (left panels) and *Fgfr3-creER*; *R26R*^tdTomato^ (right panels) femurs, pulsed at E12.5, undergone surgery at P21 and analyzed at P28. Metaphyseal (upper panels) and diaphyseal (lower panels) marrow space after one week of surgery. Grey: DIC. Scale bar: 500µm. (k): Quantification based on sections. Percentage of Col1(2.3kb)-GFP^+^tdTomato^+^ osteoblasts per total Col1(2.3kb)-GFP^+^ osteoblasts in the metaphysis (left) and diaphysis (right). *n*=4 per each group. \**p*<0.05, two-tailed, Mann-Whitney’s *U*-test. Data are presented as mean ± s.d.

To reveal the unique molecular signature characterizing Hes1-creER^+^ cell-derived BMSCs, we performed a comparative bulk RNA-seq analysis of *Cxcl12-*GFP^high^tdTomato^+^ cells isolated from *Cxcl12^GFP/+^*; *Hes1-creER*; *R26R*^tdTomato^ and *Cxcl12^GFP/+^*; *Fgfr3-creER*; *R26R*^tdTomato^ bones with tamoxifen injection at E12.5 (Supplementary Fig.11a). A principal component analysis revealed that biological replicates of Hes1^CE^-E12.5 and Fgfr3^CE^-E12.5 Cxcl12-GFP^high^ cells grouped by lineage (Supplementary Fig.11b). Notably, 2,969 genes were differentially expressed between the two groups (DEGs) (Supplementary Fig.11c,d). Gene Ontology (GO) enrichment analysis of DEGs revealed significant enrichment of a number of GO terms related to osteoblast differentiation and mineralization, including *integrin-mediated signaling pathway* (GO:0007229), *positive regulation of osteoblast differentiation* (GO:0045669), *extracellular matrix organization* (GO:0030198) and *ossification* (GO:0001503). Particularly, genes associated with osteoblast differentiation were upregulated in Cxcl12-GFP^high^Fgfr3^CE^-E12.5-tdTomato^+^ cells, including *Pth1r*, *Spp1* and *Mmp13* (Fig.4e), indicating that BMSCs derived from Fgfr3-creER^+^ cells possess a state more biased for osteoblasts than BMSCs derived from Hes1-creER^+^ cells.

Subsequently, we further set out to investigate the osteogenic potential of these two classes of BMSCs. First, we tested the response of Hes1^CE^-E12.5 and Fgfr3^CE^-E12.5 BMSCs to bone anabolic agents by giving once daily injections of PTH (1-34, 100µg/kg b.w.) to *Col1a1(2.3kb)-GFP*; *Hes1-creER* or *Fgfr3-creER*; *R26R*^tdTomato^ mice (pulsed at E12.5) for 3 weeks from P21 to P42 (Fig.4f). As expected, this anabolic regimen substantially increased the trabecular bone mass particularly in the metaphysis (Fig.4g,h) (Col1-GFP^+^ cells: 372.5±29.6 and 362.0±65.0 cells/area for Hes1^CE^-E12.5 and Fgfr3^CE^-E12.5 femurs, respectively, compared to 114.8±11.6 cells/area in Vehicle-treated mice). Importantly, Fgfr3^CE^-E12.5 cells contributed to a great majority of newly-formed osteoblasts, while virtually no Hes1^CE^-E12.5 cells contributed to these cells in the metaphysis (Fig.4g,h) (GFP^+^tdTomato^+^ cells; 2.6±2.3 % and 55.1±5.0 % of total Col1-GFP^+^ cells for Hes1^CE^-E12.5 and Fgfr3^CE^-E12.5 femurs, respectively), demonstrating that Hes1-creER^+^ cell-derived BMSCs barely account for anabolic responses during the postnatal stage. Second, we investigated an injury-responsive osteogenesis capability of Hes1^CE^-E12.5 and Fgfr3^CE^-E12.5 cells using a femoral bone marrow ablation model, which induced direct differentiation of BMSCs within the marrow cavity (Fig.4i)^40^. After 7 days of marrow ablation, Hes1^CE^-E12.5 cells contributed to *de novo* bone formation exclusively in the diaphysis, while Fgfr3^CE^-E12.5 cells contributed predominantly in the metaphysis (Fig.4j,k) (GFP^+^tdTomato^+^ cells; 4.7±3.4 % and 76.0±7.9 % of total Col1-GFP^+^ cells in metaphyseal bone marrow, 53.0±8.0 % and 9.0±3.0% of total Col1-GFP^+^ cells in diaphyseal bone marrow for Hes1^CE^-E12.5 and Fgfr3^CE^-E12.5 femurs, respectively). Therefore, Hes1-creER^+^ cell-derived BMSCs contribute to osteogenesis specifically in the diaphyseal marrow space.

## Discussion

Taken together, our study provides descriptive data on the identity, location and fate of perichondrial cells, chondrocytes and their precursors in the fetal stage, based on a combination of single cell transcriptomic analyses and *in vivo* cell-lineage analyses of identified cell populations. Single cell RNA-seq analyses of the pre-cartilaginous condensation and the cartilage template reveal a contiguous nature of the fetal chondrocyte-perichondrial cell lineage that largely shares an overlapping set of marker genes. Subsequent studies identified three previously unreported tamoxifen-inducible *creER* lines, namely *Hes1-creER* and *Dlx5-creER* that are highly active in fetal perichondrial cells, and *Fgfr3-creER* that is highly specific to fetal chondrocytes. These newly identified *creER* lines have advantages over existing lines as detailed below, therefore might be useful for designing future studies on the fetal perichondrium.

First, we identified that *Hes1-creER* allows marking an early perichondrial population of skeletal progenitor cells in a SOX9-negative domain of the condensation. *Prrx1-cre* and *Prrx1-creER* are widely utilized tools that mark skeletal progenitor cells at the condensation stage; however, the caveat of these transgenes is that they simultaneously mark SOX9^+^ cells within the condensation. However, we acknowledge that *Hes1-creER* has limitations, in that it also marks non-skeletogenic cell populations in the limb bud. It may be advantageous to utilize multiple tools in tandem to compliment the strengths and weaknesses of each line.

Importantly, our findings from *Hes1-creER*-based lineage-tracing experiments may pose a challenge the established concept that all osteo-chondroprogenitors are derived from Sox9^+^ (Sox9-cre^+^) cells. There are two scenarios explaining the difference. First, it is possible that *Hes1-creER* marks the precursors for Sox9^+^ (Sox9-cre^+^) cells, supporting the notion that Hes1-creER^+^ cells function as earlier perichondrial skeletal progenitor cells. Second, it is also possible that *Hes1-creER* marks perichondrial cells after these cells have turned off *Sox9* (*Sox9-cre*) expression. Identifying the precise mechanism will be important to define the origin and the function of early perichondrial cells in endochondral development.

Second, we identified that *Dlx5-creER* represents a highly perichondrial cell-specific genetic tool, which provides substantial technological and conceptual advances in this study. *Dlx5-creER*-based lineage-tracing experiments unambiguously demonstrate the existence of a self-renewing population of skeletal progenitor cells in the fetal perichondrium, which provide a continuous source of precursors for Osx-creER^+^ perichondrial cells.

Third, we identified that *Fgfr3-creER* is highly specific to chondrocytes, with a significant advantage over previously reported *Acan-creER* and *Col2a1-creER* lines. *Acan* is expressed in the perichondrium; as a result, *Acan-creER* marks a large number of perichondrial cells, at least in perinatal mice^12^. In addition, we showed in this study that *Col2a1-creER* marks a substantial number of fetal perichondrial cells. In contrast, *Fgfr3-creER* marks almost exclusively chondrocytes within the cartilage template, marking little in the perichondrium. Therefore, this highly specific *Fgfr3-creER* may be useful for analyzing chondrocytes in the fetal stage.

Further, identification of *Dlx5-creER* and *Fgfr3-creER* as highly specific inducible genetic tools for perichondrial cells and chondrocyte, respectively, allows us to demonstrate that these two types of cells contribute to the functionally distinct bone marrow stromal compartments in postnatal mice. However, we acknowledge that, as *Fgfr3-creER* can mark a small number of perichondrial cells, it cannot be entirely excluded that bone marrow stromal cells are also derived from Fgfr3-creER^+^ perichondrial cells, not only from Fgfr3-creER^+^ chondrocytes.

In addition, we acknowledge that the conclusions of this study may be valid for developing femurs, but not for other skeletal elements that develop earlier or later than the femur during the fetal life in mice. A different time point for tamoxifen injection would be necessary to correctly analyze cell fates in another skeletal element.

A combination of multiple inducible mouse genetics tools in this study raises an intriguing possibility that fetal perichondrial cells might provide a complementary source of skeletal progenitor cells through a distinct route during formation of the marrow space (see the proposed diagram in Fig.5). Fetal perichondrial cells might contribute particularly to important stages of early endochondral bone development when the demand for cytogenesis is especially high; first, when the mesenchymal condensation expands explosively, and second, when the nascent marrow space is established within the cartilage template. In contrast, in later stages of development, skeletal progenitor cells established within the cartilage template appear to be self-sufficient to sustain cytogenesis for continual endochondral bone growth due to their robust self-renewal capacity; these cells also give rise to PTHrP^+^ chondrocytes with skeletal stem cell properties within the resting zone of the postnatal growth plate^36^.

**Figure 5.**
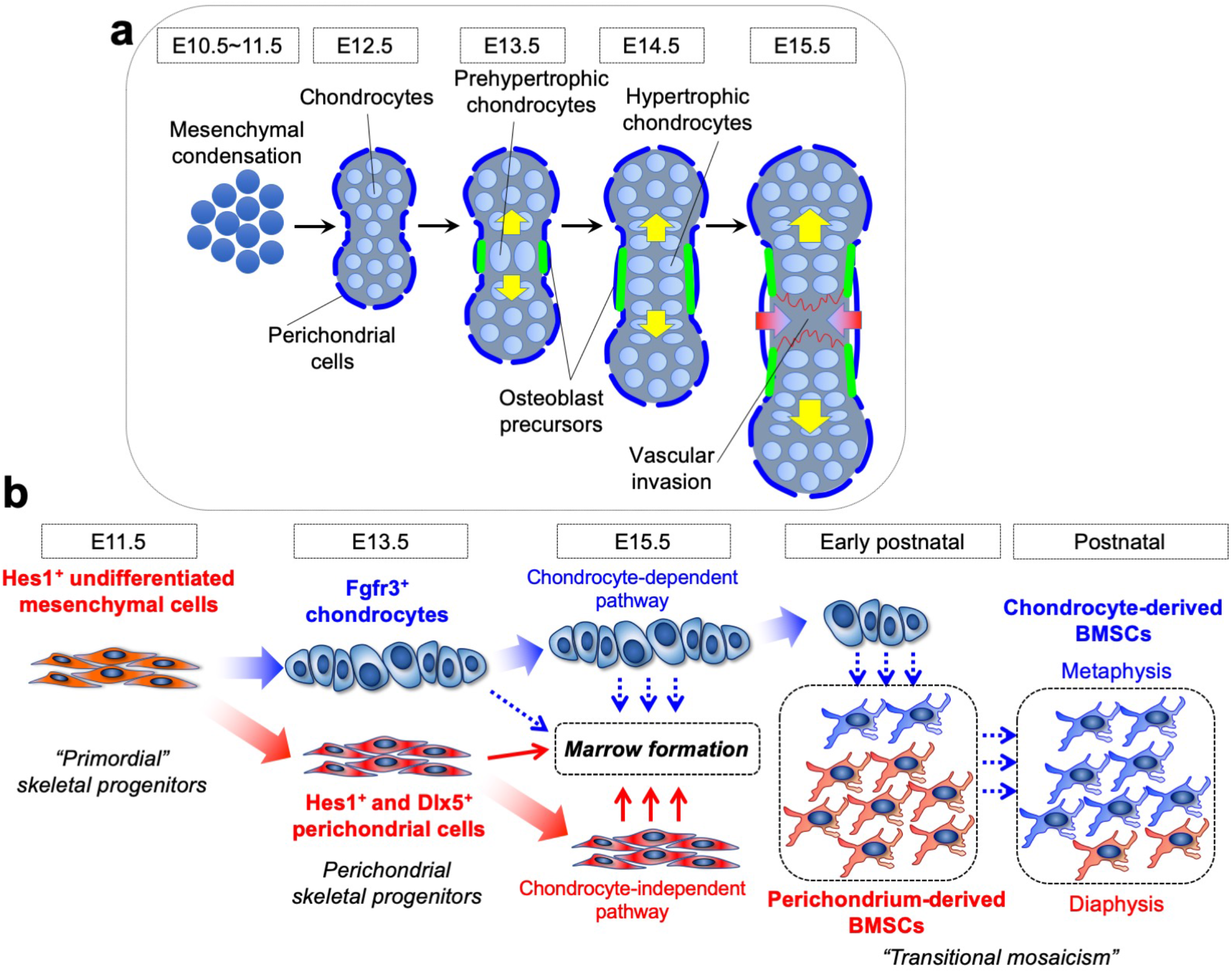
Notch effector Hes1 marks an early perichondrial population of skeletal progenitor cells in endochondral bone development. **(a)** Process of endochondral bone development and bone marrow formation. Skeletal primordium is formed by condensing mesenchymal cells. These mesenchymal cells differentiate into chondrocytes and their surrounding perichondrial cells. Both cartilage template and perichondrium contribute to endochondral bone development by providing skeletal progenitor cells. These cells in concert form the nascent marrow space along with invading endothelial and hematopoietic cells. **(b)** Early perichondrial population of skeletal progenitor cells in endochondral bone development. Undifferentiated mesenchymal cells surrounding condensation or cartilage template provide an important source during fetal endochondral bone development. During the condensation stage, Hes1^+^ undifferentiated mesenchymal cells take two distinct routes to participate in endochondral bone development. First, these cells translocate into the cartilage template and directly differentiate into Sox9^+^ chondrocytes, and then contribute to bone formation (chondrocyte-dependent pathway). Second, these cells contribute to the outer layer of the perichondrium and become Hes1^+^ and Dlx5^+^ perichondrial cells. These Hes1^+^ and Dlx5^+^ perichondrial cells translocate into the nascent marrow space and directly differentiate into marrow mesenchymal cells by bypassing a Sox9^+^ state (chondrocyte-independent pathway). These perichondrial skeletal progenitor cells keep providing osteoblasts and bone marrow stromal cells even during the postnatal stage, unlike Osx^+^ perichondrial osteoblast precursors with limited life span. As a result, postnatal bone narrow is characterized by transitional mosaicism composed of both chondrocyte-derived and perichondrium-derived stromal cells.

The adult periosteum is derived from the fetal perichondrium. In fact, adult periosteal cells exhibit greater clonogenic and regenerative potential than bone marrow stromal cells, although they share the common embryonic origin and specific markers^10^. Adult periosteal cells might maintain some of the key features of fetal perichondrial cells, such as the ability to generate chondrocytes and skeletal progenitor cells to direct endochondral ossification. How cells marked by *creER* lines identified in this study contribute to fracture healing in adult bones need to be defined through further investigation.

Importantly, the fetal perichondrial route appears to be transient, rendering the bone marrow compartment transitional in terms of the developmental origin during postnatal development. While the colony-forming fraction of the early postnatal bone marrow compartment is largely contributed by Hes1-creER^+^ cells, cells contributed by Fgfr3-creER^+^ cells become predominant in later stages in the newly formed portion of endochondral bones in the metaphysis. These latter cells appear to be the major contributor of new bone formation in response to bone anabolic agents. This unique partnership between the perichondrium and the cartilage template may be one of the major driving forces for explosive mammalian endochondral bone development.

## Acknowledgement

We thank T. Nagasawa (Osaka University) for *Cxcl12^GFP/+^* mice, C. Murtaugh *for Hes1-creER* mice, and H.M. Kronenberg for *Osx-creER* mice. This research was supported by National Institute of Health grants R01DE026666 (to N.O.), R03DE027421 (to W.O.), R01AR065409 (to S.W.) and University of Michigan MCubed 2.0 Grant (to N.O., W.O. and S.W.), MCubed 3.0 Grant (to N.O., W.O. and J.D.W.), JSPS Overseas Research Fellowship and JSPS KAKENHI Grant JP19K19236 to Y.M. We thank M. Pihalja and K. Saiya-Cork of the University of Michigan Flow Cytometry Core, Yusuke Ono of Kumamoto University, Masaki Matsushita of Nagoya University and T. Lau for supporting this study.

## Author contributions

Y.M. and N.O. conceived the project and wrote the manuscript. Y.M., M.N., W.O. and N.O. performed the experiment. J.D.W. performed the bioinformatic analysis. S.Y.W. provided the mice and critiqued the manuscript.

## Methods

### Mouse strains

*Osx-creER*^6^, *Hes1-creER*^21^ and *Cxcl12^GFP/+^*^41^ mice have been described previously. *Prrx1-cre* (JAX005584), *Col2a1-cre* (JAX003554), *Col2a1-creER* (JAX006774), *Fgfr3-creER* (JAX025809), *Dlx5-creER* (JAX010705), *Acta1-cre* (JAX006139), *Myl-cre* (JAX024713), *Mck-cre* (JAX006475), *Tie2-cre* (JAX008863), *Rosa26-CAG-loxP-stop-loxP-tdTomato* (Ai14: *R26R-tdTomato*, JAX007914), *Rosa26-SA-loxP-GFP-stop-loxP-DTA* (JAX006331), *Col1a1(2.3kb)-GFP* (JAX013134), *Osteocalcin-GFP* (JAX017469) and *CBF:H2B-Venus* (JAX020942) mice were acquired from the Jackson laboratory. *Fgfr3-GFP* (MMRRC:012012-UCD) mice were acquired from the Mutant Mouse Resource and Research Centers. All procedures were conducted in compliance with the Guidelines for the Care and Use of Laboratory Animals approved by the University of Michigan’s Institutional Animal Care and Use Committee (IACUC), protocol 7681 (Ono). All mice were housed in a specific pathogen-free condition, and analyzed in a mixed background. Mice were identified by micro-tattooing or ear tags. Tail biopsies of mice were lysed by a HotShot protocol (incubating the tail sample at 95°C for 30 min in an alkaline lysis reagent followed by neutralization) and used for PCR-based genotyping (GoTaq Green Master Mix, Promega, and Nexus X2, Eppendorf). Perinatal mice were also genotyped fluorescently (BLS miner’s lamp) whenever possible. Mice were euthanized by over-dosage of carbon dioxide or decapitation under inhalation anesthesia in a drop jar (Fluriso, Isoflurane USP, VetOne).

### Tamoxifen and induction of *cre-loxP* recombination

Tamoxifen (Sigma T5648) and 4-hydroxytamoxifen (4-OHT, Sigma H6278) were mixed with 100% ethanol until completely dissolved. Subsequently, a proper volume of sunflower seed oil (Sigma S5007) was added to the tamoxifen-ethanol mixture and rigorously mixed. The tamoxifen-ethanol-oil mixture was incubated at 60°C in a chemical hood until the ethanol evaporated completely. The tamoxifen-oil mixture was stored at room temperature until use. Tamoxifen was injected intraperitoneally using a 26-1/2-gauge needle (BD309597).

### Intermittent administration of parathyroid hormone (PTH)

Mice were given once daily subcutaneous injections of PTH (1-34, 100µg/kg b.w., 4095855, Bachem) for 3 weeks from P21 to P42.

### Bone marrow ablation surgery

Mice were anesthetized through a nose cone, in which 1.5-2% isoflurane was constantly provided with oxygen through a vaporizer. For bone marrow ablation surgery, right femurs were operated, while left femurs were untreated and used as an internal control. After skin incision, cruciate ligaments of the knee were carefully separated using a dental excavator, and a hole was made in the intercondylar region of femurs using a 26-1/2-gauge needle (BD309597). The endodontic instruments (K-file, #35, 40, 45, 50) (GC, Tokyo, Japan) were used in a stepwise manner to remove a cylindrical area of the marrow space. The surgical field was irrigated with saline, and the incision line was sutured.

### Histology and immunohistochemistry

Samples were dissected under a stereomicroscope (Nikon SMZ-800), and fixed in 4% paraformaldehyde for a proper period, typically ranging from 3 hours to overnight at 4°C, then decalcified in 15% EDTA for a proper period, typically ranging from 3 hours to 14 days. Embryonic samples were not decalcified. Subsequently, samples were cryoprotected in 30% sucrose/PBS solutions and then in 30% sucrose/PBS:OCT (1:1) solutions, each at least overnight at 4°C. Samples were embedded in an OCT compound (Tissue-Tek, Sakura) under a stereomicroscope and transferred on a sheet of dry ice to solidify the compound. Embedded samples were cryosectioned at 14µm using a cryostat (Leica CM1850) and adhered to positively charged glass slides (Fisherbrand ColorFrost Plus). Sections were postfixed in 4% paraformaldehyde for 15 min at room temperature. For immunostaining, sections were permeabilized with 0.25% TritonX/TBS for 30 min, blocked with 3% BSA/TBST for 30 min and incubated with rabbit anti-Sox9 polyclonal antibody (1:500, EMD-Millipore, AB5535), rat anti-endomucin (Emcn) monoclonal antibody (1:100, Santa Cruz Biotechnology, sc65495), rabbit anti-Myh3 polyclonal antibody 1:500, Abcam, ab124205 or rabbit anti-Osx polyclonal antibody (1:500, Abcam, ab22552) overnight at 4°C, and subsequently with Alexa Fluor 647-conjugated donkey anti-rabbit IgG (A31573) or Alexa Fluor 633-conjugated goat anti-rat IgG (A21049) (1:400, Invitrogen) for 3 hours at room temperature. Sections were further incubated with DAPI (4’,6-diamidino-2-phenylindole, 5µg/ml, Invitrogen D1306) to stain nuclei prior to imaging.

### RNAscope in situ hybridization

Samples were fixed in 4% paraformaldehyde overnight at 4°C, and then cryoprotected. Frozen sections at 14µm were prepared on positively charged glass slides. In situ hybridization was performed with RNAscope 2.5 HD Reagent kit Brown (Advanced Cell Diagnostics 322300) using following probes: *Hes1* (417701), *Fgfr3* (440771) and *Sox9* (custom-designed) according to the manufacturer’s protocol.

### Imaging and cell quantification

Images were captured by an automated inverted fluorescence microscope with a structured illumination system (Zeiss Axio Observer Z1 with ApoTome.2 system) and Zen 2 (blue edition) software. The filter settings used were: FL Filter Set 34 (Ex. 390/22, Em. 460/50 nm), Set 38 HE (Ex. 470/40, Em. 525/50 nm), Set 43 HE (Ex. 550/25, Em. 605/70 nm), Set 50 (Ex. 640/30, Em. 690/50 nm) and Set 63 HE (Ex. 572/25, Em. 629/62 nm). The objectives used were: Fluar 2.5x/0.12, EC Plan-Neofluar 5x/0.16, Plan-Apochromat 10x/0.45, EC Plan-Neofluar 20x/0.50, EC Plan-Neofluar 40x/0.75, Plan-Apochromat 63x/1.40. Images were typically tile-scanned with a motorized stage, Z-stacked and reconstructed by a maximum intensity projection (MIP) function. Differential interference contrast (DIC) was used for objectives higher than 10x. Representative images of at least three independent biological samples are shown in the figures. Quantification of cells on sections was performed using NIH Image J software.

### Cell preparation

For embryonic samples, hind limbs were harvested and incubated with 2 Wunsch units of Liberase TM (Sigma/Roche 5401127001) in 2ml Ca^2+^, Mg^2+^-free Hank’s Balanced Salt Solution (HBSS, Sigma H6648) at 37°C for 15 min on a shaking incubator (ThermomixerR, Eppendorf). For postnatal samples, soft tissues and epiphyses were carefully removed from dissected femurs. After removing distal epiphyseal growth plates and cutting off proximal ends, femurs were cut roughly and incubated with 2 Wunsch units of Liberase TM and 1mg of Pronase (Sigma/Roche 10165921001) in 2ml Ca^2+^, Mg^2+^-free HBSS at 37°C for 60 min on a shaking incubator. After cell dissociation, cells were mechanically triturated using an 18-gauge needle with a 1ml Luer-Lok syringe (BD) and a pestle with a mortar (Coors Tek), and subsequently filtered through a 70µm cell strainer (BD) into a 50ml tube on ice to prepare single cell suspension. These steps were repeated for 5 times, and dissociated cells were collected in the same tube. Cells were pelleted and resuspended in an appropriate medium for subsequent purposes. For cell culture experiments, cells were resuspended in 10ml culture medium and counted on a hemocytometer.

### Flow cytometry

Dissociated cells were stained by standard protocols with the following antibodies (1:500, eBioscience). eFluor450-conjugated CD31 (390, endothelial/platelet), CD45 (30F-11, hematopoietic), Ter119 (TER-119, erythrocytes), allophycocyanin (APC)-conjugated CD31 (390, endothelial/platelet), CD45 (30F-11, hematopoietic) and Ter119 (TER-119, erythrocytes). Flow cytometry analysis was performed using a four-laser BD LSR Fortessa (Ex. 405/488/561/640 nm) and FACSDiva software. Acquired raw data were further analyzed on FlowJo software (TreeStar). Representative plots of at least three independent biological samples are shown in the figures.

### Single cell RNA-seq analysis of fluorescence-activated cell sorting (FACS)-isolated cells

Cell sorting was performed using a four-laser BD FACS Aria III (Ex.407/488/561/640nm) high-speed cell sorter with a 100µm nozzle. Cells of interest (tdTomato^+^ cells for the experiment described in Fig.1 and Supplementary Fig.1, *Cxcl12*-GFP^high^ and/or tdTomato^+^ [both single and double positive: GFP^high^, tdTomato^+^ and GFP^high^tdTomato^+^] cells for the experiment of Fig.3 and Supplementary Fig.6) were directly sorted into ice-cold DPBS/1% BSA, pelleted by centrifugation and resuspended in appropriate amount of DPBS/1% BSA (1,000 cells/μl). Cell numbers were quantified by Countless II automated Cell Counter (ThermoFisher) before loading onto the Chromium Single Cell 3’ v2 microfluidics chip (10x Genomics Inc., Pleasanton, CA). cDNA libraries were sequenced by Illumina HiSeq 4000 using two lanes and 50 cycle paired-end read, generating a total of ∼ 770 million reads, or by Illumina or NovaSeq 6000. The sequencing data was first pre-processed using the 10X Genomics software Cell Ranger. For alignment purposes, we generated and used a custom genome fasta and index file by including the sequences of *eGFP* and *tdTomato-WPRE* to the mouse genome (mm10). Further downstream analysis steps were performed using the Seurat^19^ and Monocle^25^ R package. We filtered out cells with less than 500 genes per cell and with more than 20% mitochondrial read content. The downstream analysis steps include normalization, identification of highly variable genes across the single cells, scaling based on number of UMI, dimensionality reduction (PCA, CCA and t-SNE), unsupervised clustering, and the discovery of differentially expressed cell-type specific markers. Differential gene expression to identify cell-type specific genes was performed using the nonparametric Wilcoxon rank sum test.

To integrate *Hes1* and *Fgfr3* scRNA-seq data, we used LIGER^42^. This method performs integrative nonnegative matrix factorization (iNMF) to identify shared and dataset-specific metagene factors, which produces an aligned latent space. LIGER then uses this aligned space for joint clustering and *t*-SNE visualization. For our analysis here, we used *k* = 25 factors and default clustering parameters.

### Code availability

LIGER algorithm is fully available as described^42^.

### Bulk RNA-seq analysis of FACS-isolated cells

Dissociated bone marrow cells harvested from P21 littermate mice were pooled based on the genotype [*Cxcl12^GFP/+^*; *Hes1-creER*; *R26R*^tdTomato^ (pulsed at E12.5, Hes1-E12.5 mice) or *Cxcl12^GFP/+^*; *Fgfr3-creER*; *R26R*^tdTomato^ (pulsed at E12.5, Fgfr3-E12.5 mice)]. Cell sorting was performed using a four-laser BD FACS Aria III (Ex.407/488/561/640nm) high-speed cell sorter with a 100µm nozzle. *Cxcl12*-GFP^high^tdTomato^+^ cells isolated from Hes1-E12.5 or Fgfr3-E12.5 mice were directly sorted into ice-cold DPBS/10% FBS and pelleted by centrifugation. Total RNA was isolated using PicoPure RNA Isolation Kit (KIT0204, ThermoFisher), followed by DNA-free DNA removal kit (AM1906, ThermoFisher) to remove contaminating genomic DNA. RNA Integrity Number (RIN) was assessed by Agilent 2100 Bioanalyzer RNA 6000 Pico Kit. Samples with RIN>8.0 were used for subsequent analyses. Complementary DNAs were prepared by SMART-Seq v4 Ultra Low Input RNA Kit for Sequencing (Clontech 634888). Post-amplification quality control was performed by Agilent TapeStation DNA High Sensitivity D1000 Screen Tape system. DNA libraries were prepared by Nextera XT DNA Library Preparation Kit (Illumina) and submitted for NextGen sequencing (Illumina HiSeq 4000). DNA libraries were sequenced using following conditions; six samples per lane, 50 cycle single-end read. Reads files were downloaded and concatenated into a single .fastq file for each sample. The quality of the raw reads data for each sample was checked using FastQC to identify quality problems. Tuxedo Suite software package was subsequently used for alignment (using TopHat and Bowtie2), differential expression analysis, and post-analysis diagnostics. FastQC was used for a second round of quality control (post-alignment). HTSeq/DESeq2 was used for differential expression analysis using UCSC mm10 as the reference genome sequence. Only non-ambiguously mapped reads were counted. Meta-analysis of RNA-seq data was performed using iPathwayGuide software (Advaita).

### Colony-forming assay and subcloning

Nucleated bone marrow cells were plated into tissue culture 6 well plates (BD Falcon) at a density of <10^5^ cells/cm^2^, and cultured in low-glucose DMEM with GlutaMAX supplement (Gibco 10567022) and 10% mesenchymal stem cell-qualified FBS (Gibco 12662029) containing penicillin-streptomycin (Sigma P0781) for 10∼14 days. Cell cultures were maintained at 37°C in a 5% CO_2_ incubator. Representative images of at least three independent biological samples are shown in the figures. For CFU-Fs, cells were fixed with 70% Ethanol for 5 min and stained for 2% methylene blue.

Colonies marked by tdTomato were individually subcloned using a method described previously^36^. Briefly, a tile-scanned virtual reference of tdTomato^+^ colonies and cloning cylinders (Bel-Art) were used to isolate an individual colony. Single cell-derived clones of tdTomato^+^ cells were cultured in a basal medium described above at 37°C with 5% CO_2_ with exchanges into fresh media every 3–4 days.

### Statistical analysis

Results are presented as mean values ± S.D. Statistical evaluation was conducted using the Mann-Whitney’s *U*-test or one-way ANOVA. A P value of <0.05 was considered significant. No statistical method was used to predetermine sample size. Sample size was determined on the basis of previous literature and our previous experience to give sufficient standard deviations of the mean so as not to miss a biologically important difference between groups. The experiments were not randomized. All of the available mice of the desired genotypes were used for experiments. The investigators were not blinded during experiments and outcome assessment. One femur from each mouse was arbitrarily chosen for histological analysis. Genotypes were not particularly highlighted during quantification.

### Data availability

The datasets generated during and/or analyzed during the current study are available from the corresponding author on reasonable request. The single cell and bulk RNA-seq datasets presented herein have been deposited in the National Center for Biotechnology Information (NCBI)’s Gene Expression Omnibus (GEO), and are accessible through GEO Series accession numbers GSE126966 [https://www.ncbi.nlm.nih.gov/geo/query/acc.cgi?acc=GSE126966], GSE126967 [https://www.ncbi.nlm.nih.gov/geo/query/acc.cgi?acc=GSE126967], GSE126968 [https://www.ncbi.nlm.nih.gov/geo/query/acc.cgi?acc=GSE126968], GSE126969 [https://www.ncbi.nlm.nih.gov/geo/query/acc.cgi?acc=GSE126969] and GSE144411 [https://www.ncbi.nlm.nih.gov/geo/query/acc.cgi?acc=GSE144411]. The source data underlying all Figures and Supplementary Figures are provided as a Source Data file.

---

**Supplementary Figure 1.**
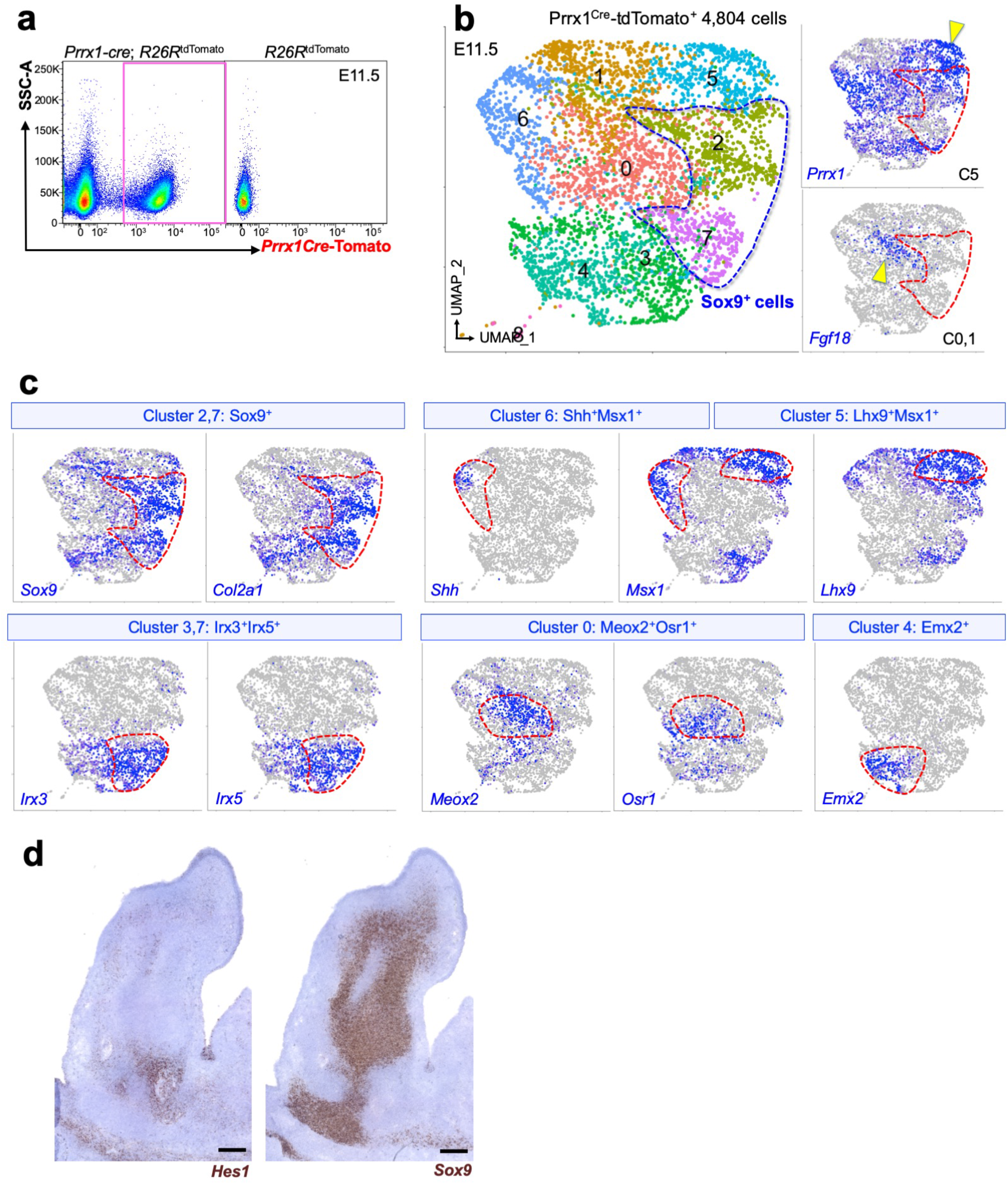
Single cell RNA-seq analysis of limb bud mesenchymal cells. **(a-c)** Single cell RNA-seq analysis of *Prrx1-cre*-marked limb bud mesenchymal cells at E11.5. (a): FACS-sorting strategy for Prrx1^cre^-tdTomato^+^ cells (red box), cells isolated from *Prrx1-cre*; *R26R*^tdTomato^ limbs. Right panel: control cells. (b): UMAP-based visualization of major classes of Prrx1^cre^-tdTomato^+^ cells (left) and feature plots of representative markers (right). Blue: high expression. Dotted contour: Sox9^+^ cells. *n*=4,804 cells. Pooled from *n*=5 mice. (c): Feature plots of genes enriched in each cluster. Cluster 2,7: *Sox9*^+^, Cluster 6: *Shh*^+^*Msx1*^+^, Cluster 5: *Lhx9*^+^*Msx1*^+^, Cluster 3,7: *Irx3*^+^*Irx5*^+^, Cluster 0: *Meox2*^+^*Osr1*^+^, Cluster 4: *Emx2*^+^. Red contour: featured clusters. **(d)** RNAscope in situ hybridization analysis of E11.5 limb bud for *Hes1* and *Sox9* mRNA. Scale bar: 200µm. *n*=3 mice.

**Supplementary Figure 2.**
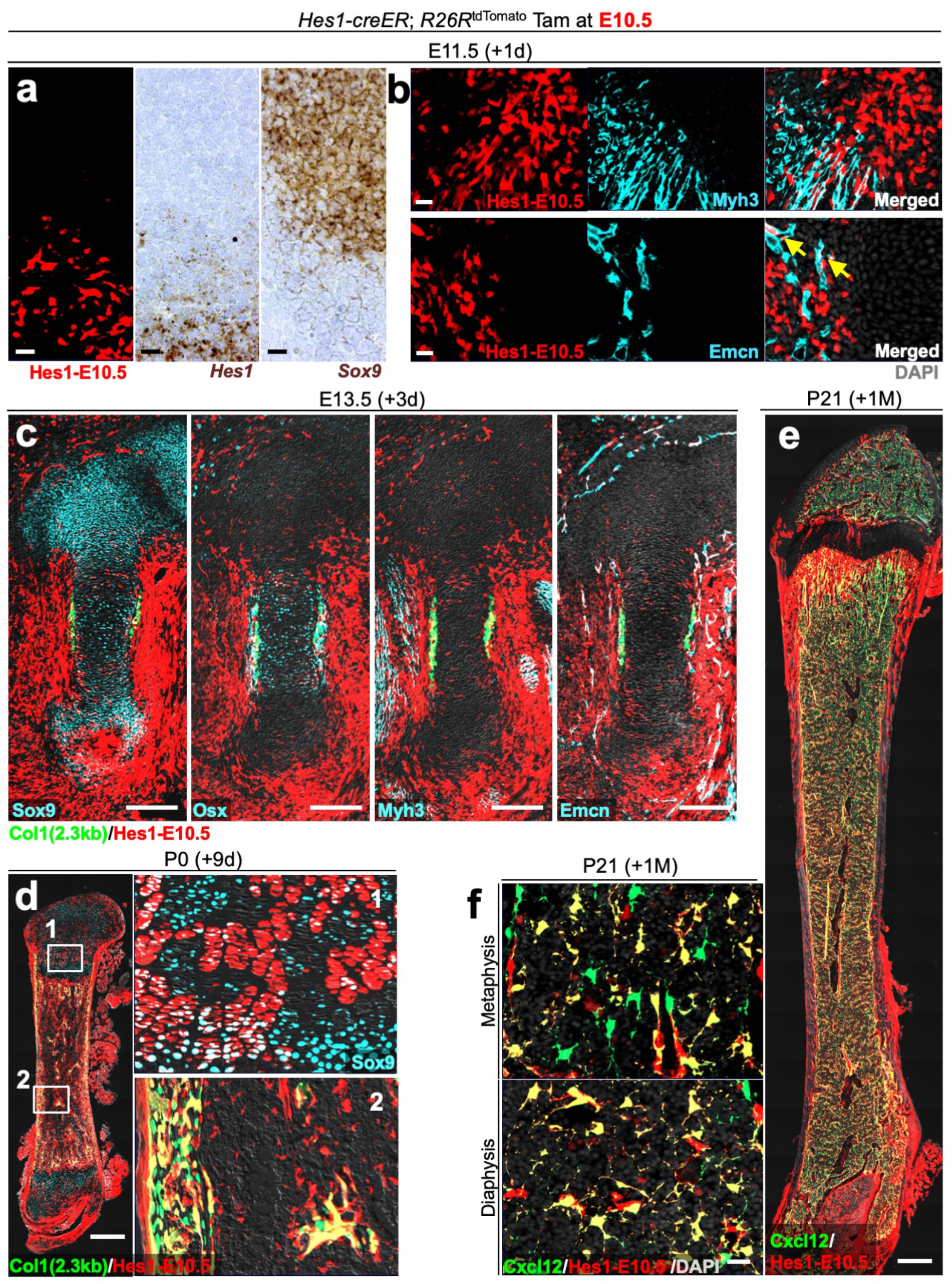
Contribution of Hes1^+^Sox9^neg^ mesenchymal cells to endochondral bone development. Cell-fate analysis of *Hes1-creER*^+^ mesenchymal cells of the condensation stage, pulsed at E10.5. *Hes1-creER*; *R26R*^tdTomato^ femurs carrying *Col1a1(2.3kb)-GFP* (c,d) or *Cxcl12-GFP* (e,f) reporters. (a): Short-chase analysis of *Hes1-creER^+^* cells at E11.5. Mesenchymal condensation. RNAscope in situ hybridization analysis for *Hes1* (center) and *Sox9* (right) mRNA. Scale bar: 20µm. (b): Immunostaining for Myh3 (top) and Emcn (bottom). Arrow: *Hes1-creER*^+^Emcn^+^ cells. Grey: DAPI. Scale bar: 20µm. (c): Cartilage template at E13.5, after 3 days of chase. Imunostaining for Sox9, Osx, Myh3 and Emcn. Grey: DIC. Scale bar: 200µm. (d): Neonatal femur at P0, after 9 days of chase. Right panels: magnified views of (1: growth plate, 2: bone marrow). Grey: DIC. Scale bar: 200µm. (e): Whole femur at P21 with growth plates on top, after one month of chase. Grey: DIC. Scale bar: 500µm. (f): metaphyseal bone marrow (top) and diaphyseal bone marrow (bottom). Grey: DAPI. Scale bar: 20µm. *n*=4 mice per each time point.

**Supplementary Figure 3.**
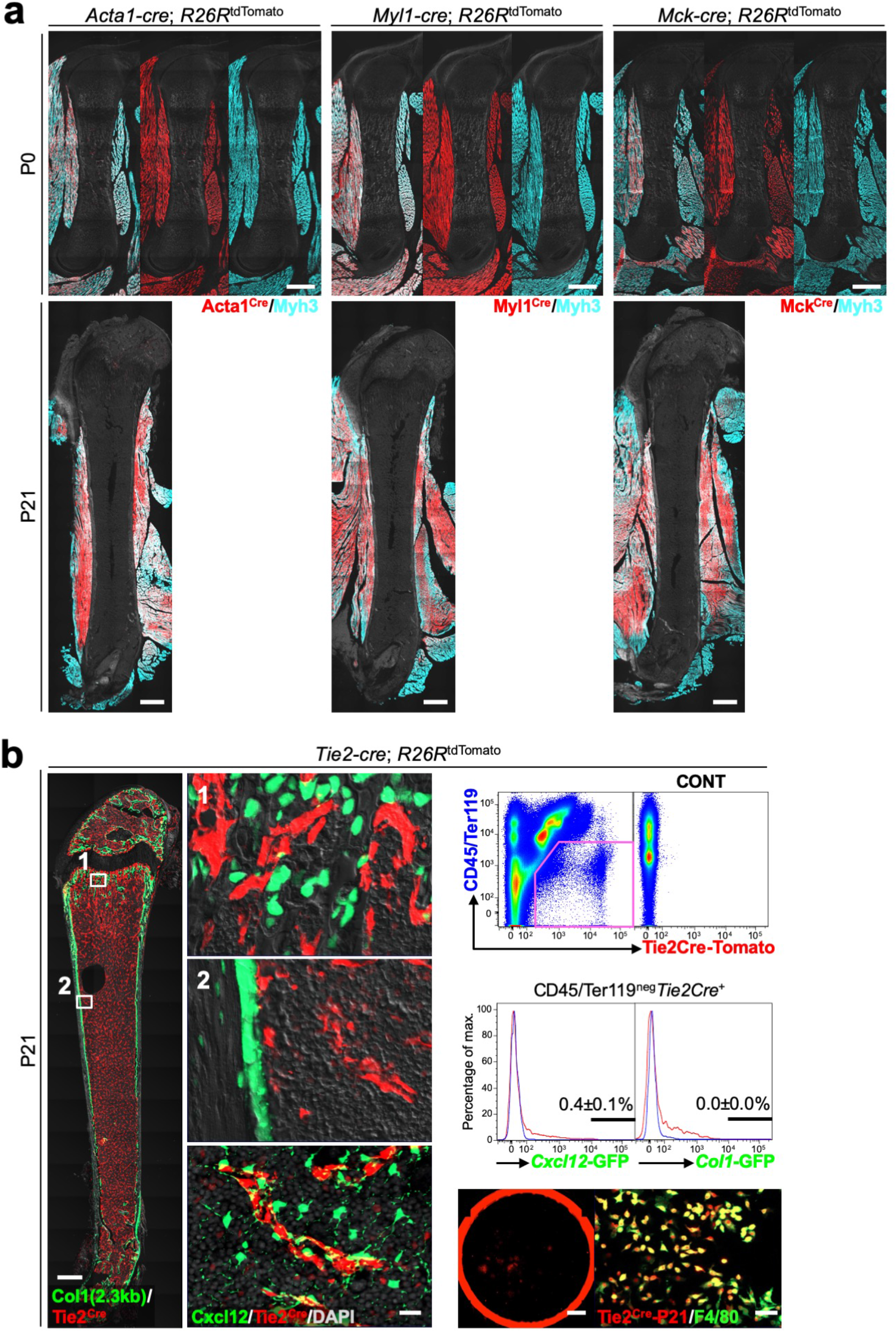
Skeletal muscle cells or endothelial cells do not contribute to chondrocytes, osteoblasts or bone marrow stromal cells. **(a)** Fate mapping analysis of *Acta1-cre, Myl1-cre or Mck-cre*-marked skeletal muscle cells at P0 (top) and P21 (bottom). *Acta1-cre*; *R26R*^tdTomato^ (left), *Myl1-cre*; *R26R*^tdTomato^ (center) and *Mck-cre*; *R26R*^tdTomato^ (right) femurs. Immunostaining for Myh3. Grey: DIC. Scale bar: 500µm. *n*=3 mice per each group. **(b)** Fate-mapping analysis of *Tie2-cre*-marked endothelial cells at P21. *Col1a1(2.3kb)-GFP*; *Tie2-cre*; *R26R*^tdTomato^ and *Cxcl12^GFP/+^*; *Tie2-cre*; *R26R*^tdTomato^ femurs. Left panel: whole bone. Grey: DIC. Scale bar: 500µm. Upper center panels: magnified views of boxed areas (1: trabecular bone, 2: endosteal marrow space). Lower center panel: magnified view of diaphyseal bone marrow of *Cxcl12^GFP/+^*; *Tie2-cre*; *R26R*^tdTomato^ femur. Grey: DAPI. Scale bar: 20µm. *n*=3 mice per each group. Upper right panels: flow cytometry analysis of bone marrow cells. Right center panel: histogram showing GFP expression. Blue lines: control cells. *n*=5 (*Cxcl12^GFP/+^*), *n*=4 (*Col1a1(2.3kb)-GFP*) mice. Lower right panels: CFU-F assay of *Tie2-cre*; *R26R*^tdTomato^ bone marrow cells. Left: tdTomato epifluorescence. Scale bar: 5mm. Right: F4/80 immunostaining. Scale bar: 50µm. *n*=3 mice.

**Supplementary Figure 4.**
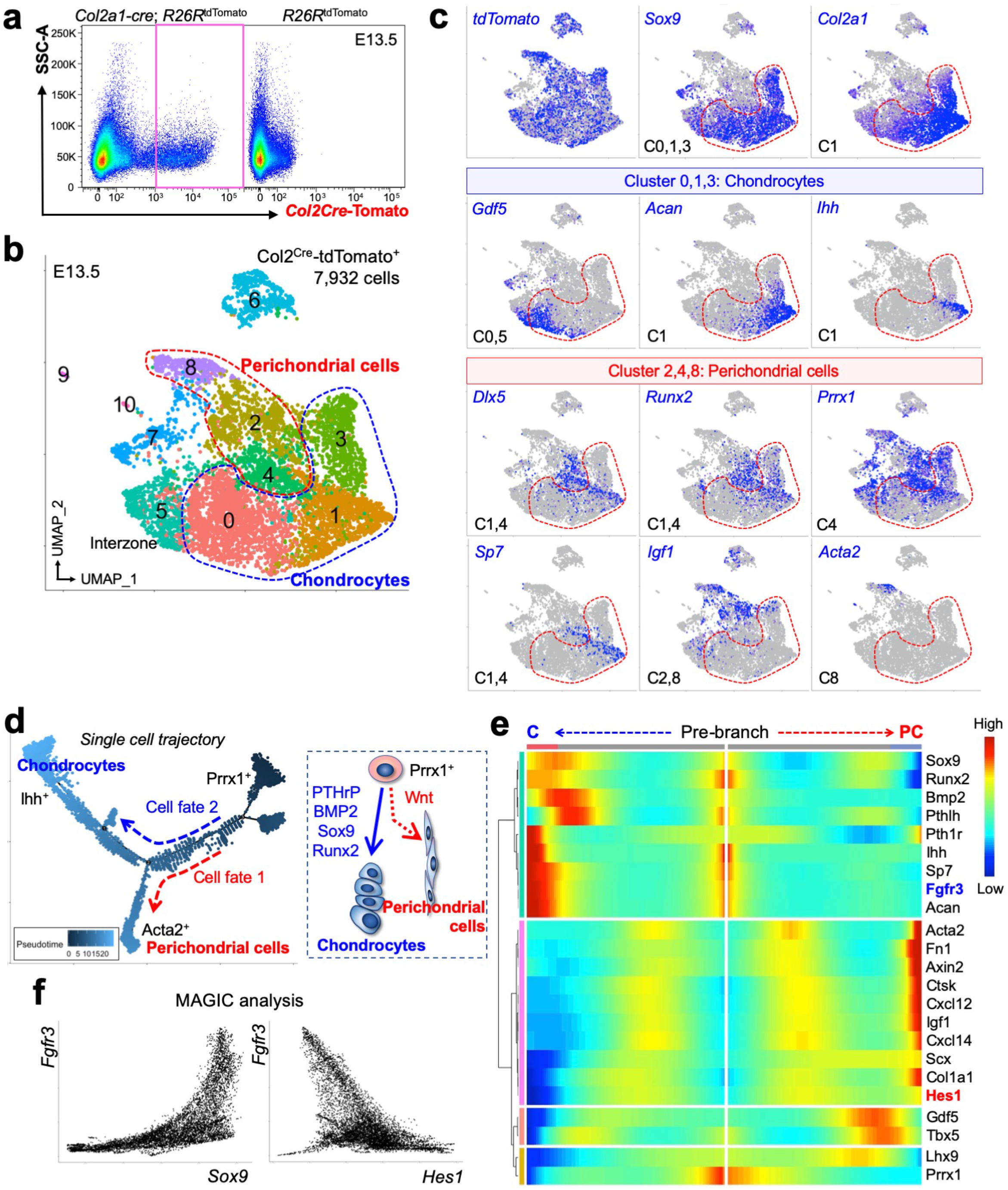
Single cell RNA-seq analysis of fetal chondrocytes and perichondrial cells. Single cell RNA-seq analysis of *Col2a1-cre*-marked chondrocytes and perichondrial cells at E13.5. (a): FACS-sorting strategy for Col2a1^cre^-tdTomato^+^ cells (red box), cells isolated from *Col2a1-cre*; *R26R*^tdTomato^ limbs. Right panel: control cells. (b): UMAP-based visualization of major classes of Col2a1^cre^-tdTomato^+^ cells. Red contour: perichondrial cells, Blue contour: chondrocytes. *n*=7,932 cells. Pooled from *n*=5 mice. (c): Feature plots of genes enriched in chondrocyte clusters and perichondrial cell clusters. Blue: high expression. Cluster 0, 1, 3: chondrocytes, cluster 2,4,8: perichondrium. Dotted contour: chondrocytes. (d): Pseudotime and single cell trajectory analysis by Monocle. Right panel: inferred lineage trajectory. (e): Heatmap of representative pseudotime-dependent differentially expressed genes. (f): MAGIC imputation analysis showing the *Sox9*-*Fgfr3* and *Hes1*-*Fgfr3* relationships.

**Supplementary Figure 5.**
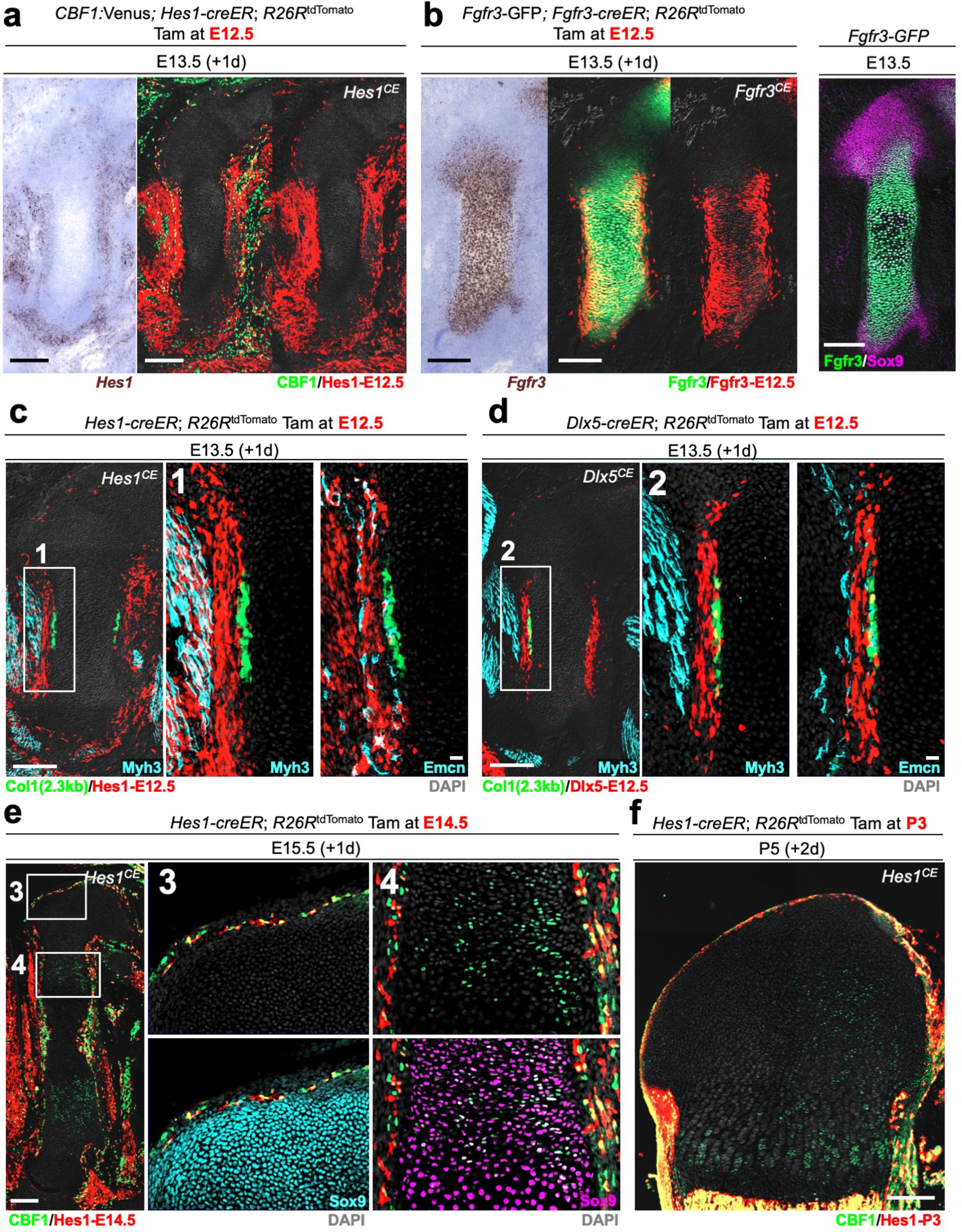
*Hes1-creER* marks self-renewing Dlx5^+^Osx^neg^ progenitor cells in the fetal perichondrium. **(a-d)** Cell-fate analysis of *Hes1-creER*^+^, *Dlx5-creER*^+^ or *Fgfr3-creER*^+^ cells, pulsed at E12.5. (a): *CBF1:Venus*; *Hes1-creER*; *R26R*^tdTomato^ femurs at E13.5. Left: RNAscope in situ hybridization analysis of *Hes1* mRNA. Grey: DIC. Scale bar: 200µm. (b): *Fgfr3*-GFP; *Fgfr3-creER*; *R26R*^tdTomato^ femurs at E13.5. Left: RNAscope in situ hybridization analysis of *Fgfr3* mRNA. Right: *Fgfr3*-GFP femur at E13.5 with Sox9 immunostaining. Grey: DIC. Scale bar: 200µm. (c): *Col1a1(2.3kb)-GFP*; *Hes1-creER*; *R26R*^tdTomato^ femurs at E13.5. Left: Myh3 immunostaining. Grey: DIC. Scale bar: 200µm. Center: magnified views of boxed area (1). Grey: DAPI. Right: Emcn immunostaining. Grey: DAPI. Scale bar: 20µm. (d) *Col1a1(2.3kb)-GFP*; *Dlx5-creER*; *R26R*^tdTomato^ femurs at E13.5. Left: Myh3 immunostaining. Grey: DIC. Scale bar: 200µm. Center: magnified views of boxed area (2). Grey: DAPI. Right: Emcn immunostaining. Grey: DAPI. Scale bar: 20µm. *n*=4 mice per each group. **(e)** *CBF1:Venus*; *Hes1-creER*; *R26R*^tdTomato^ femurs at E15.5, pulsed at E14.5. Left: Grey: DIC. Scale bar: 200µm. Center, right: magnified views of boxed area (3: articular cartilage, 4: proliferating chondrocytes) with Sox9 immunostaining. Grey: DAPI. *n*=4 mice. **(f)** *CBF1:Venus*; *Hes1-creER*; *R26R*^tdTomato^ femurs at P5, pulsed at P3. Grey: DIC. Scale bar: 200µm. *n*=4 mice.

**Supplementary Figure 6.**
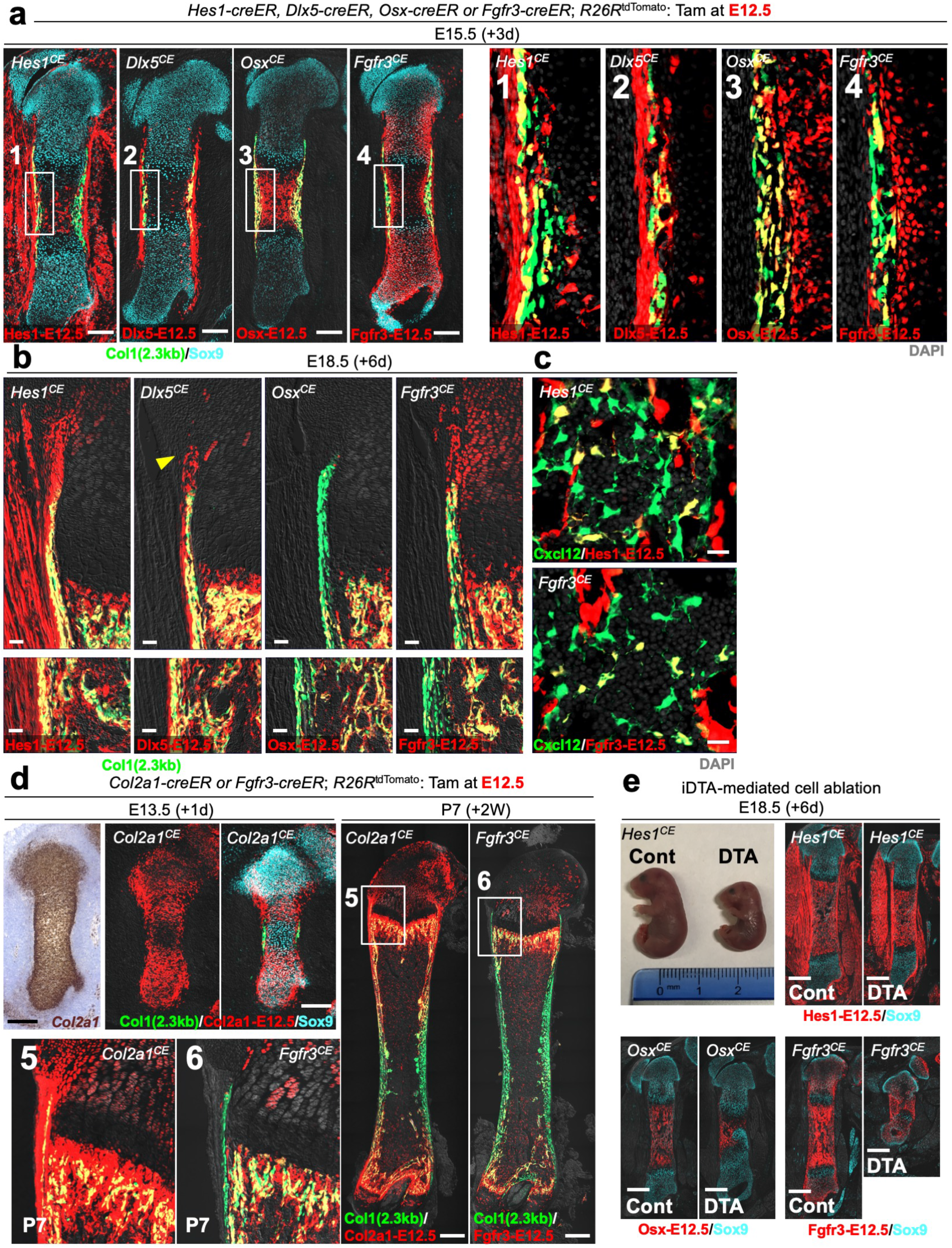
*Hes1-creER* marks self-renewing Dlx5^+^Osx^neg^ progenitor cells in the fetal perichondrium. **(a-c)** Cell-fate analysis of *Hes1-creER*^+^, *Dlx5-creER*^+^, *Osx-creER*^+^ or *Fgfr3-creER*^+^ cells, pulsed at E12.5, carrying *Col1a1(2.3kb)-GFP* (a,b) or *Cxcl12-GFP* (c) reporters. (a): Left panels: whole femurs at E15.5. Grey: DIC. Scale bar: 200µm. Right panels: magnified view of (1-4). Grey: DAPI. *n*=4 mice per each group. (b): Perichondrium, bone collar and growth plate at E18.5. Arrowhead: groove of Ranvier. Grey: DIC. Scale bar: 50µm. *n*=4 mice per each group. (c) Bone marrow at P0. Grey: DAPI. Scale bar: 20µm. *n*=3 mice per each group. **(d)** Cell-fate analysis of *Col2a1-creER*^+^ or *Fgfr3-creER*^+^ cells, pulsed at E12.5. Upper left: *Col1a1(2.3kb)-GFP*; *Col2a1creER*; *R26R*^tdTomato^ femurs at E13.5. Grey: DIC. Scale bar: 200µm. *n*=3 mice. Right: *Col1a1(2.3kb)-GFP*; *Col2a1-creER*; *R26R*^tdTomato^ or *Col1a1(2.3kb)-GFP*; *Fgfr3-creER*; *R26R*^tdTomato^ femurs at P7. Scale bar: 500µm. Lower left: magnified view of (5,6: perichondrium, bone collar and growth plate). Grey: DIC. *n*=3 mice per each group. **(e)** Inducible partial cell ablation of *Hes1-creER*^+^, *Osx-creER*^+^ or *Fgfr3-creER*^+^ cells using an inducible Diphtheria toxin fragment A (iDTA) allele. *Hes1-creER*, *Osx-creER or Fgfr3-creER*; *Rosa26*^lsl-tdTomato/+^ (Cont) and *Hes1-creER*, *Osx-creER or Fgfr3-creER*; *Rosa26*^lsl-tdTomato/iDTA^ (DTA) mice at E18.5, pulsed at E12.5. *n*=10 (Hes1-Cont), 7 (Hes1-DTA), 6 (Fgfr3-DTA), 5 (Osx-Cont), 4 (Osx-DTA, Fgfr3-Cont).

**Supplementary Figure 7.**
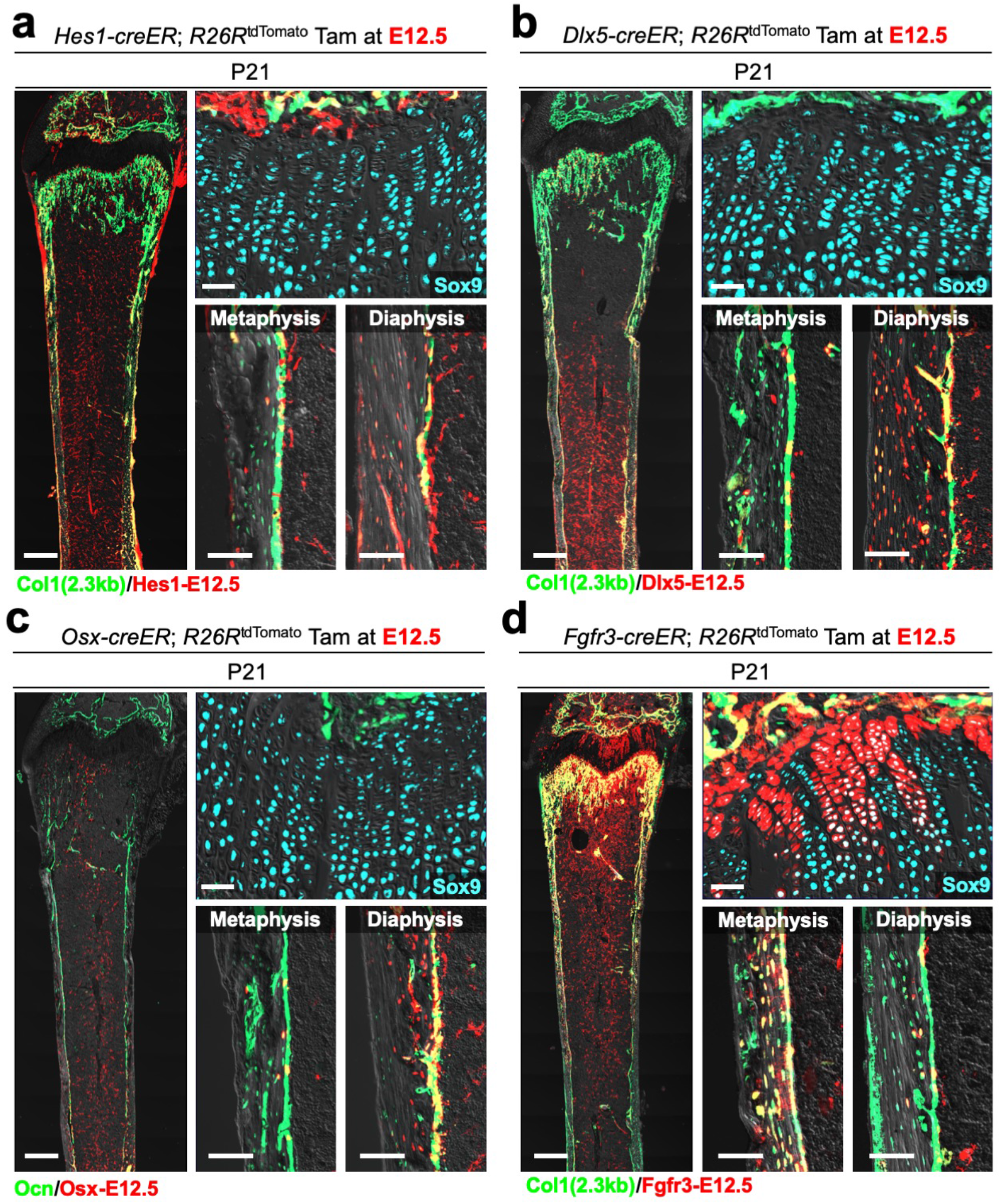
Differential contribution of Hes1^+^, Dlx5^+^, Osx^+^ and Fgfr3^+^ cells to postnatal growth plate and bone marrow stromal compartment. **(a-d)** Contribution of fetal *Hes1-creER*^+^, *Dlx5-creER*^+^, *Osx-creER*^+^ or *Fgfr3-creER*^+^ cells (pulsed at E12.5) to growth plate chondrocytes and osteoblasts at P21. *Col1a1(2.3kb)-GFP*; *Hes1-creER*; *R26R*^tdTomato^ (a), *Col1a1(2.3kb)-GFP*; *Dlx5-creER*; *R26R*^tdTomato^ (b), *Osteocalcin (Ocn)*; *Osx-creER*; *R26R*^tdTomato^ (c) or *Col1a1(2.3kb)-GFP*; *Fgfr3-creER*; *R26R*^tdTomato^ femurs (d) with growth plates on top. Whole bone (left), growth plates (upper right), endocortical marrow space at metaphysis (lower center) and diaphysis (lower right). Grey: DIC (a-d). Scale bar: 500µm (left panels), 50µm (upper right panels), 100µm (lower center and right panels). *n*=3 mice per each group.

**Supplementary Figure 8.**
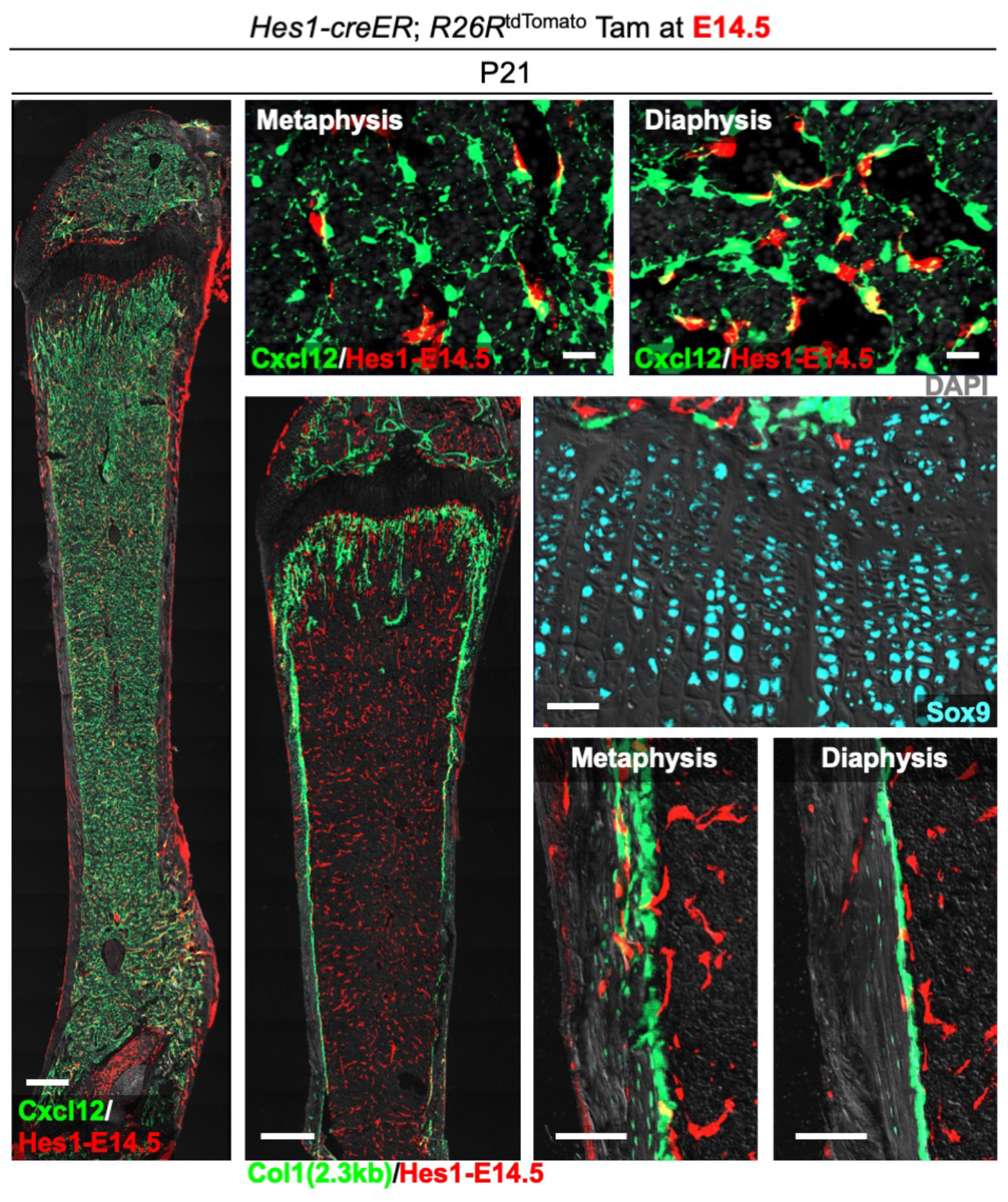
*Hes1-creER* predominantly marks endothelial cells in late fetal stages. Contribution of *Hes1-creER*^+^ cells (pulsed at E14.5) to Cxcl12-GFP^+^ bone marrow stromal cells, growth plate chondrocytes or osteoblasts at P21. *Cxcl12^GFP/+^*; *Hes1-creER*; *R26R^tdTomato^* (left, upper) femurs with growth plates on top. Left: Whole bone. Grey: DIC. Scale bar: 500µm. Upper: Metaphyseal and diaphyseal bone marrow. Grey: DAPI. Scale bar: 20µm. *Col1a1(2.3kb)-GFP*; *Hes1-creER*; *R26R^tdTomato^* (lower center, center right, lower right) femurs with growth plates on top. Whole bone (lower center), growth plates (center right), endocortical marrow space at metaphysis and diaphysis (lower right). Grey: DIC. Scale bar: 500µm (lower center), 50µm (center right), 100µm (lower right). *n*=3 mice per each group.

**Supplementary Figure 9.**
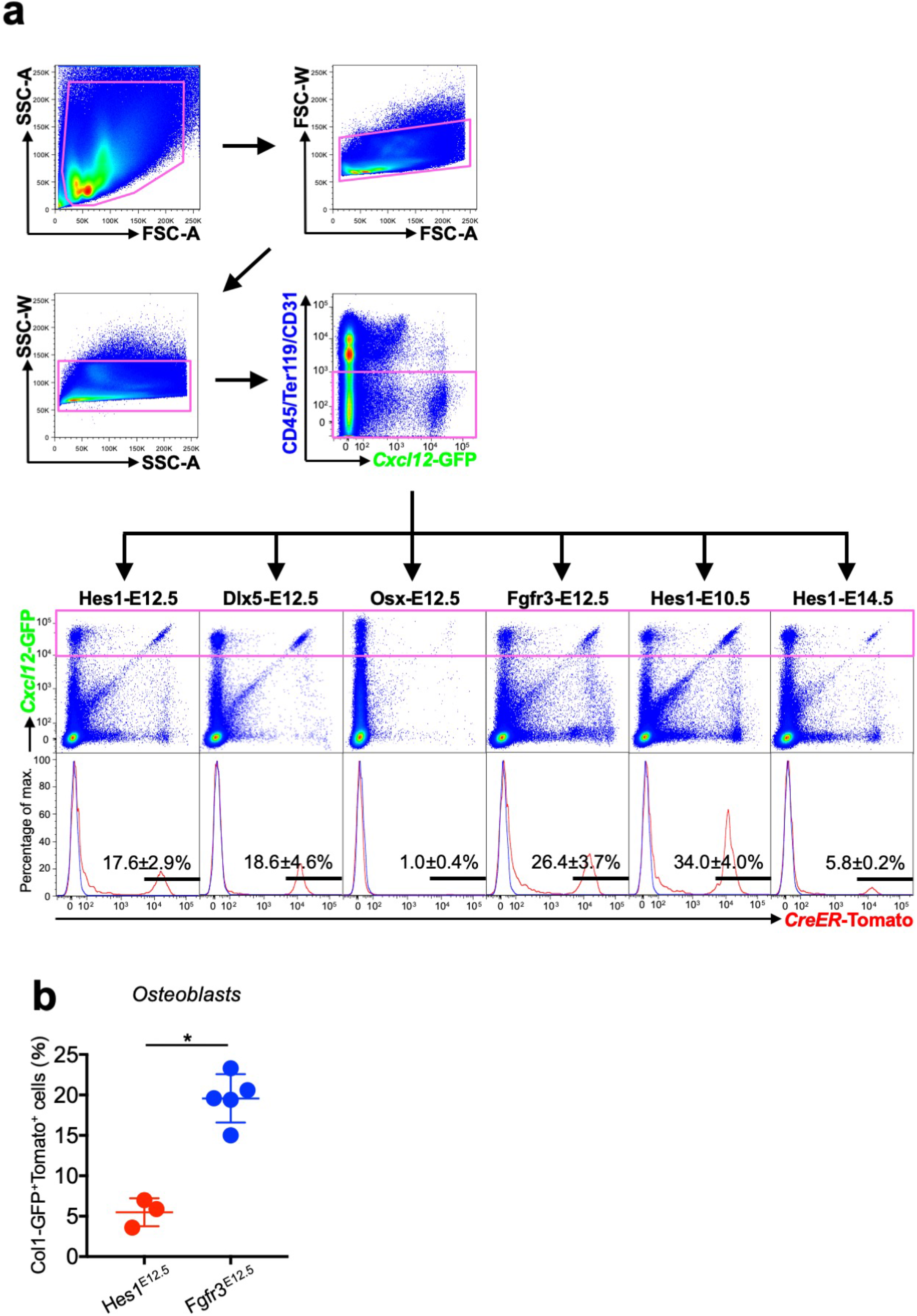
Flow cytometry analysis of lineage-marked Cxcl12-GFP^+^ bone marrow stromal cells. **(a)** Flow cytometry analysis of CD45/Ter119/ CD31^neg^ cells at P21. Gating strategy for bone marrow cells isolated from *Cxcl12^GFP/+^*; *Hes1-creER*; *R26R*^tdTomato^ femurs (pulsed at E10.5, E12.5 or E14.5), *Cxcl12^GFP/+^*; *Dlx5-creER*; *R26R*^tdTomato^ femurs (pulsed at E12.5), *Cxcl12^GFP/+^*; *Osx-creER*; *R26R*^tdTomato^ femurs (pulsed at E12.5) and *Cxcl12^GFP/+^*; *Fgfr3-creER*; *R26R*^tdTomato^ femurs (pulsed at E12.5). *n*=5 (Hes1-E12.5), *n*=4 (Dlx5-E12.5), *n*=5 (Osx-E12.5), *n*=10 (Fgfr3-E12.5), *n*=6 (Hes1-E10.5), *n*=4 (Hes1-E14.5) mice per group. **(b)** Percentage of Col1(2.3kb)-GFP^+^tdTomato^+^ osteoblasts per total Col1(2.3kb)-GFP^+^ osteoblasts. *n*=3 (Hes1-E12.5), *n*=5 (Fgfr3-E12.5) mice. \**p*<0.05, two-tailed, Mann-Whitney’s *U*-test. Data are presented as mean ± s.d.

**Supplementary Figure 10.**
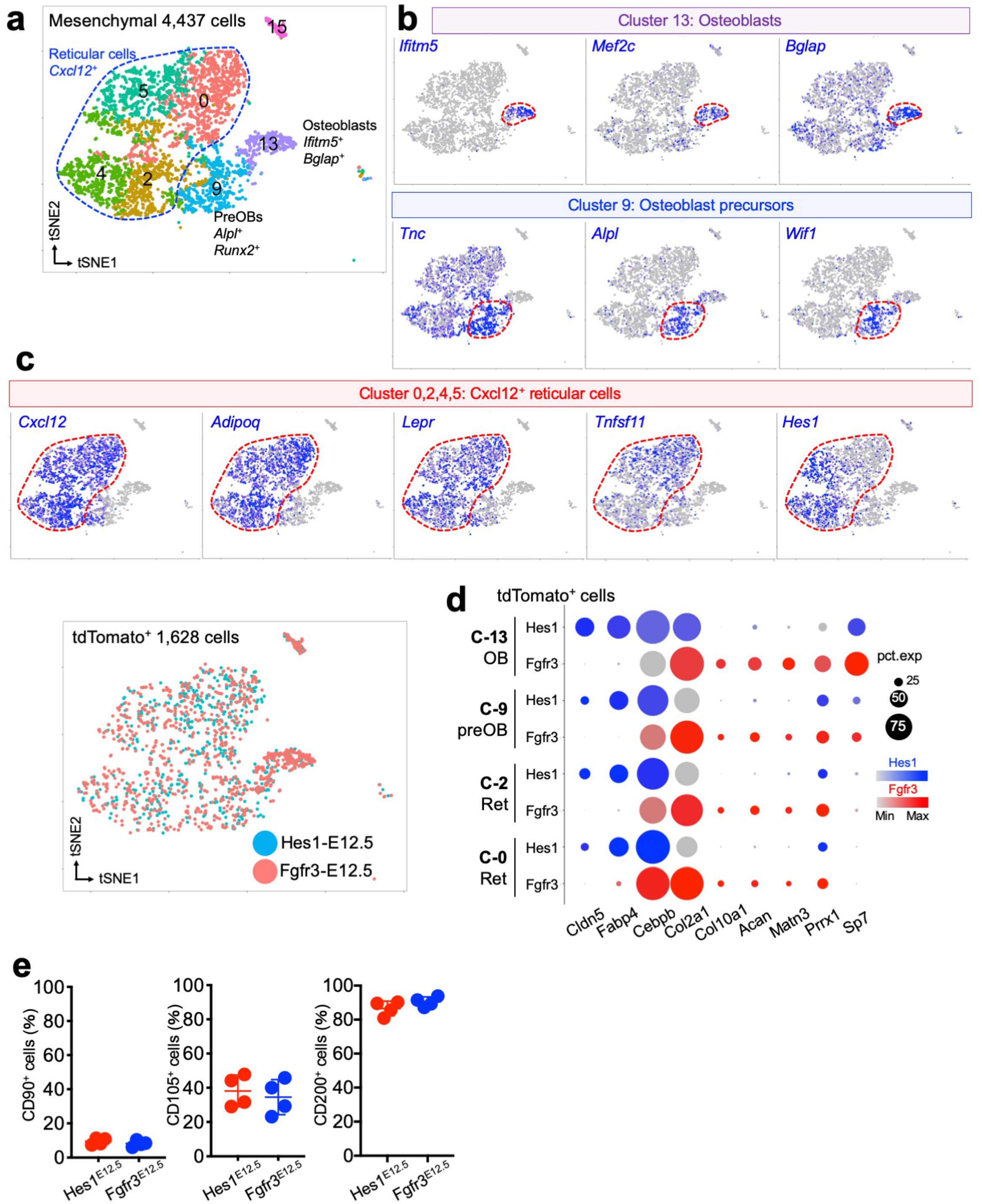
Combined lineage-tracing and single cell RNA-seq analyses of bone marrow stromal cells derived from fetal perichondrial cells and chondrocytes. **(a-d)** Combined lineage-tracing and single cell RNA-seq analyses of Cxcl12-GFP^+^ bone marrow stromal cells and tdTomato^+^ mesenchymal cells derived from fetal Hes1^+^ perichondrial cells and Fgfr3^+^ chondrocytes. Both single and double positive cells (GFP^high^, tdTomato^+^ and GFP^high^tdTomato^+^) were FACS-isolated from *Cxcl12^GFP/+^*; *Hes1-creER*; *R26R*^tdTomato^ and *Cxcl12^GFP/+^*; *Fgfr3-creER*; *R26R*^tdTomato^ femurs at P21, pulsed at E12.5 (Hes1-E12.5 or Fgfr3-E12.5). Two single cell datasets were integrated by LIGER. (a): *t*-SNE-based visualization of major classes of mesenchymal Cxcl12-GFP^high^ and/or tdTomato^+^ (Hes1-E12.5 or Fgfr3-E12.5) cells (Cluster 0, 2, 4, 5, 9, 13, 15). *n*=4,437 cells. Pooled from *n*=4 mice per group. (b,c): Feature plots. Cluster 13: Osteoblasts. Cluster 9: Osteoblast precursors. Cluster 0, 2, 4, 5: *Cxcl12*^+^ reticular cells. Blue: high expression. (d): Split-dot-based visualization of representative gene expression. Circle size: percentage of cells expressing a given gene in a given cluster (0 – 100%), Color density: expression level of a given gene. **(e)** Flow cytometry-based analysis of Cxcl12-GFP^high^tdTomato^+^ cells (Hes1-E12.5 or Fgfr3-E12.5). Percentage of CD90^+^, CD105^+^ and CD200^+^ cells per total Cxcl12-GFP^high^tdTomato^+^ cells. *n*=4 per group. Two-tailed, Mann-Whitney’s *U*-test. Data are presented as mean ± s.d.

**Supplementary Figure 11.**
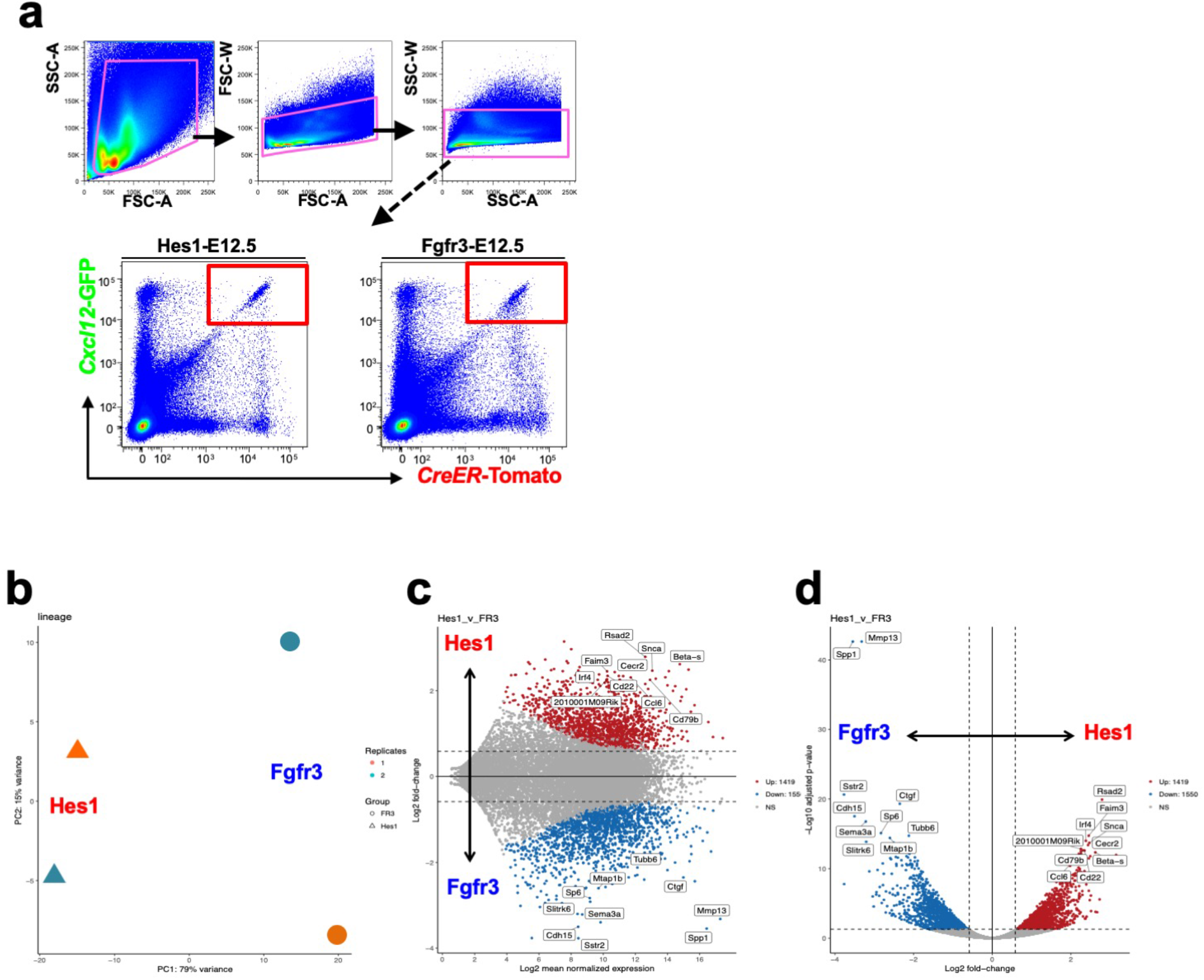
Comparative RNA-seq analysis of bone marrow stromal cells derived from fetal perichondrial cells and chondrocytes. **(a-c)** Comparative bulk RNA-seq analysis of Cxcl12-GFP^high^ stromal cells derived from fetal Hes1^+^ perichondrial cells and Fgfr3 ^+^ chondrocytes. (a): FACS-sorting of Cxcl12-GFP^high^tdTomato^+^ (Hes1-E12.5 and Fgfr3-E12.5) cells. Gating strategy for bone marrow cells isolated from *Cxcl12^GFP/+^*; *Hes1-creER*; *R26R*^tdTomato^ and *Cxcl12^GFP/+^*; *Fgfr3-creER*; *R26R*^tdTomato^ mice, pulsed at E12.5. (b): Principal component analysis (PCA) plot of four biological samples. Triangles: Cxcl12-GFP^high^Hes1-E12.5 cells, circles: Cxcl12-GFP^high^Fgfr3-E12.5 cells. *x*-axis: PC1, 79% variance, *y*-axis: PC2, 15% variance. *n*=2 biological replicates (Hes1-E12.5: pooled from *n*=4 mice, Fgfr3-E12.5: pooled from *n*=5 mice). (c): MA plot, *x*-axis: log_2_FPKM, *y*-axis: log_2_fold-change. y > 0.58 represents genes enriched in Cxcl12-GFP^high^Hes1-E12.5 (red), y < − 0.58 represents genes enriched in Cxcl12-GFP^high^Fgfr3-E12.5 (blue). (d): Volcano plot, *x*-axis: log_2_fold-change, *y*-axis: −log_10_adjusted *p*-value. x > 0.58 represents genes enriched in Cxcl12-GFP^high^Hes1-E12.5 (red), x < − 0.58 represents genes enriched in Cxcl12-GFP^high^Fgfr3-E12.5 (blue). Each dot represents a gene.

